# The centromere localization domain of kinetoplastid kinetochore protein KKT2 recognizes the free N-terminus of histone H3

**DOI:** 10.64898/2026.08.13.744621

**Authors:** Aleksandra Ciszek, Patryk Ludzia, Gabriele Marcianò, William Allen, Midori Ishii, Sam Forsyth, Christopher W Wood, Christina Redfield, Bungo Akiyoshi

## Abstract

Kinetochores are multiprotein complexes that drive chromosome segregation in eukaryotes. Kinetoplastids, a group of early-diverging eukaryotes that lack canonical kinetochore components, have a unique set of kinetochore proteins. How their kinetochores are assembled specifically at centromeres in the absence of a centromere-specific histone H3 variant CENP-A remains unknown. Here, we demonstrate that the centromere localization (CL) domain of KKT2 has similarities to a ZZ domain. An invariant aspartate present in histone H3-binding ZZ domains is conserved in the KKT2 CL domain. Using nuclear magnetic resonance (NMR) spectroscopy and isothermal titration calorimetry (ITC), we show that *Trypanosoma brucei* KKT2 CL binds the N-terminus of histone H3. Strikingly, even mono-methylation of the N-terminal amino group of histone H3 abolishes the binding. Our study raises a possibility that centromere-specific kinetochore assembly in *T. brucei* is ensured by abundant N-terminal methylations of histone H3 in non-centromeric regions.

## Introduction

Kinetochores are macromolecular protein complexes that assemble onto centromeric DNA of each chromosome and govern chromosome movement by interacting with spindle microtubules during mitosis (Musacchio and Desai, 2017). There are more than 40 kinetochore proteins, many of which are widely conserved across eukaryotes (van Hooff et al., 2017). However, none of these canonical kinetochore proteins are conserved in kinetoplastids, a group of flagellated eukaryotes that are evolutionarily divergent from traditional model eukaryotes (Berriman et al., 2005; Ivens et al., 2005; El-Sayed et al., 2005). They instead have a unique set of kinetochore proteins (Akiyoshi and Gull, 2014). Regardless of kinetochore composition, it is critical that cells control where to assemble a kinetochore on a chromosome, a mechanism called kinetochore specification (Cheeseman and Desai, 2008). Many organisms have regional centromeres whose size ranges from a few kilobases to megabases and their kinetochore specification relies on a centromere-specific histone H3 variant CENP-A, which maintains its positional information by recruiting a CENP-A-specific histone chaperone (Rowley and Jansen, 2025). It remains a mystery how kinetochores assemble specifically at centromeres without CENP-A in kinetoplastids (Lowell and Cross, 2004).

Among kinetoplastids, kinetochores and centromeres are best characterized in *Trypanosoma brucei*, an experimentally-tractable parasite that causes African trypanosomiasis (Ishii and Akiyoshi, 2022). *T. brucei* has regional centromeres, which consist of AT-rich repetitive sequences of 20–120 kb with no specific DNA sequence common to all centromeres (Obado et al., 2007; Echeverry et al., 2012; Akiyoshi and Gull, 2014; Rabuffo et al., 2024). DNA methylation is extremely rare in *T. brucei* (Militello et al., 2008), meaning that the mechanism that marks kinetochore assembly sites via DNA hypomethylation (Altemose et al., 2022) is unlikely to be in operation. *T. brucei* has four canonical histones (H2A, H2B, H3, and H4) and four histone variants (H2AZ, H2BV, H3V, and H4V), none of which is specific to centromeres (Lowell and Cross, 2004; Lowell et al., 2005; Siegel et al., 2009). While their histone-fold domains are highly conserved, the N-terminal tails are highly divergent even for canonical histones (Figueiredo et al., 2009; Deák et al., 2023, 2026). As in other eukaryotes, numerous post-translational modifications (PTMs) have been identified on the side chains of histone proteins (Janzen et al., 2006; Mandava et al., 2007; Kraus et al., 2020; Maree et al., 2022; Ocampo et al., 2025). Some PTMs are apparently unique to trypanosomatids, including histone H2A C-terminal tail hyper-acetylation (Janzen et al., 2006; Ludzia et al., 2025). Besides side chains, PTMs also exist at the N-terminal (Nα) amino group of histones (Demetriadou et al., 2020). In *T. brucei*, Nα methylations are highly abundant for all canonical histones (Janzen et al., 2006; Mandava et al., 2007; Kraus et al., 2020). The functional relevance of these Nα methylations remains unknown.

More than 20 kinetochore proteins have been identified in *T. brucei* (Akiyoshi and Gull, 2014; Nerusheva and Akiyoshi, 2016; D’Archivio and Wickstead, 2017; Nerusheva et al., 2019). Available evidence suggests that KKT2 and KKT3 form the base of kinetoplastid kinetochores and play crucial roles in recruiting other kinetochore proteins (Marcianò et al., 2021; Ishii et al., 2022). These homologous proteins have three domains conserved among kinetoplastids: an N-terminal kinase domain, a central domain that promotes their centromere localization, and a C-terminal divergent polo box domain (Nerusheva and Akiyoshi, 2016; Marcianò et al., 2021). In addition, putative DNA-binding motifs (SPKK and AT-hook) are present in some kinetoplastids (Akiyoshi and Gull, 2014). Our previous study has revealed that the central domain contains a unique zinc-binding domain (denoted the CL domain), which is essential for centromere localization, followed by a C2H2-type zinc finger (Marcianò et al., 2021). However, it remained unknown how the CL domain specifically localizes at centromeres. Here we show that the CL domain has similarities to the ZZ domain. Using NMR and ITC, we show that the KKT2 CL domain binds a histone H3 peptide, and that this binding is abolished by modifications at the very N-terminus of the peptide. On the basis of these results, we discuss a possible mechanism for how trypanosomes determine kinetochore positions.

## Results

### KKT2/3 CL domain has similarity to histone H3-binding ZZ domain

We previously solved crystal structures of the KKT2 CL domain from *Bodo saltans* and *Perkinsela*, revealing structural conservation in these divergent kinetoplastids (Marcianò et al., 2021). AlphaFold3-based models (Abramson et al., 2024) of *T. brucei* KKT2 CL and KKT3 CL show overall similarity to these crystal structures (Figure S1), suggesting that KKT2 and KKT3 CL domains share similar structures that are highly conserved among kinetoplastids. However, structural homology searches for these X-ray structures and AlphaFold3 models using DALI (Holm, 2020) or Foldseek (van Kempen et al., 2024) failed to reveal any strong hits (Marcianò et al., 2021).

Sequence analysis identified an aspartate that is strictly conserved in the CL domain of KKT2 and KKT3 (D622 in KKT2 and D692 in KKT3 in *T. brucei*), and the corresponding D-to-E mutants abolished the kinetochore localization of these CL domains, suggesting that the conserved aspartate plays an essential role for their structure or function (Marcianò et al., 2021). Our literature search has now revealed a potential similarity to a ZZ-type zinc finger (ZZ) domain, which is known to bind the N-terminus of histone H3, SUMO, or arginylated proteins (Zhang et al., 2018a; Mi et al., 2018; Liu et al., 2020; Tencer et al., 2022). Importantly, H3-binding ZZ domains have an invariant aspartate (Mi et al., 2018). Structural comparison of KKT2 CL and p300 ZZ revealed similar overall folds (Figure 1A) but the structures are circularly permuted with N- and C-termini located in different positions within the structures. This circular permutation and the different order of Cys/His pairs that coordinate zinc ions (Figure 1B) explain why our previous structural search failed to detect the similarity.

**Figure 1.**
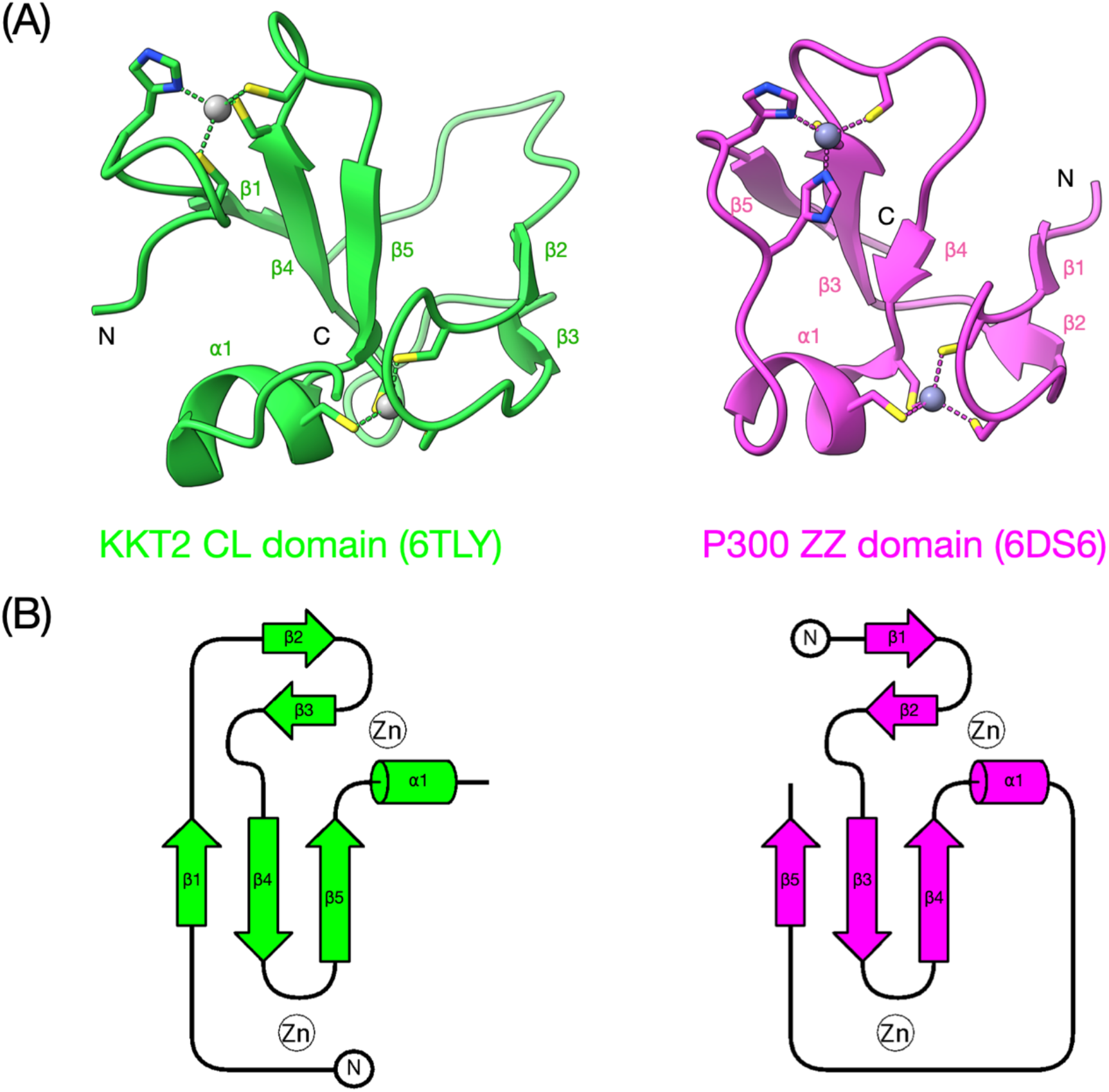
Structural similarity between KKT2 CL and ZZ domains. (A) Structures of *Bodo saltans* KKT2 CL shown in green (Protein Data Bank accession: 6TLY (Marcianò et al., 2021)) and p300 ZZ domain shown in magenta (Protein Data Bank accession: 6DS6 (Zhang et al., 2018b)). Zinc ions are shown as grey spheres. Side chains of the residues that coordinate zinc ions are shown. (B) Topology diagram of *Bodo saltans* KKT2 CL and p300 ZZ domain. Note that the first β-strand in KKT2 CL is located in the C-terminal region of the p300 ZZ domain. See also Figure S1.

### KKT2 CL domain binds the histone H3 N-terminal peptide

The structural similarity to H3-binding ZZ domains prompted us to investigate whether the *T. brucei* KKT2/3 CL domain can bind a peptide derived from the histone H3 N-terminus. In this study, we focus on the KKT2 CL domain because our attempts to purify the recombinant KKT3 CL protein in large quantities have been unsuccessful. Although histone tails are divergent in kinetoplastids (Figueiredo et al., 2009), the first eight amino acids of histone H3 in *T. brucei* are very similar to those in other eukaryotes (Figure 2). To investigate whether the KKT2 CL domain binds the N-terminal tail of histone H3, we initially employed 1D ^1^H NMR spectroscopy using unlabeled recombinant *T. brucei* KKT2^562–630^ protein (KKT2 CL hereafter) (Figure S2). The overlaid 1D spectra in Figure 3A show changes in peaks in the upfield methyl region upon addition of an N-terminal H3 12-residue peptide confirming an interaction.

**Figure 2.**
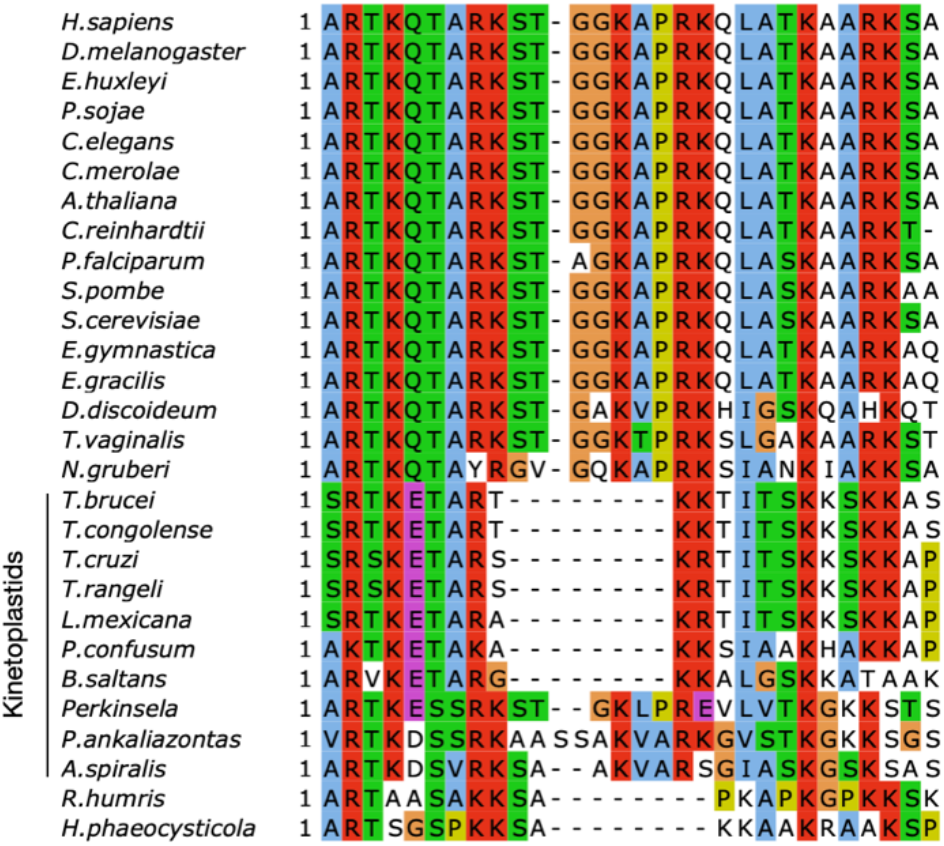
Conservation and divergence of histone H3 N-terminal tail in diverse eukaryotes. Multiple sequence alignment shows that the first eight amino acids of histone H3 in kinetoplastids are widely conserved among eukaryotes, while the rest of the tail shows a high level of divergence.

**Figure 3.**
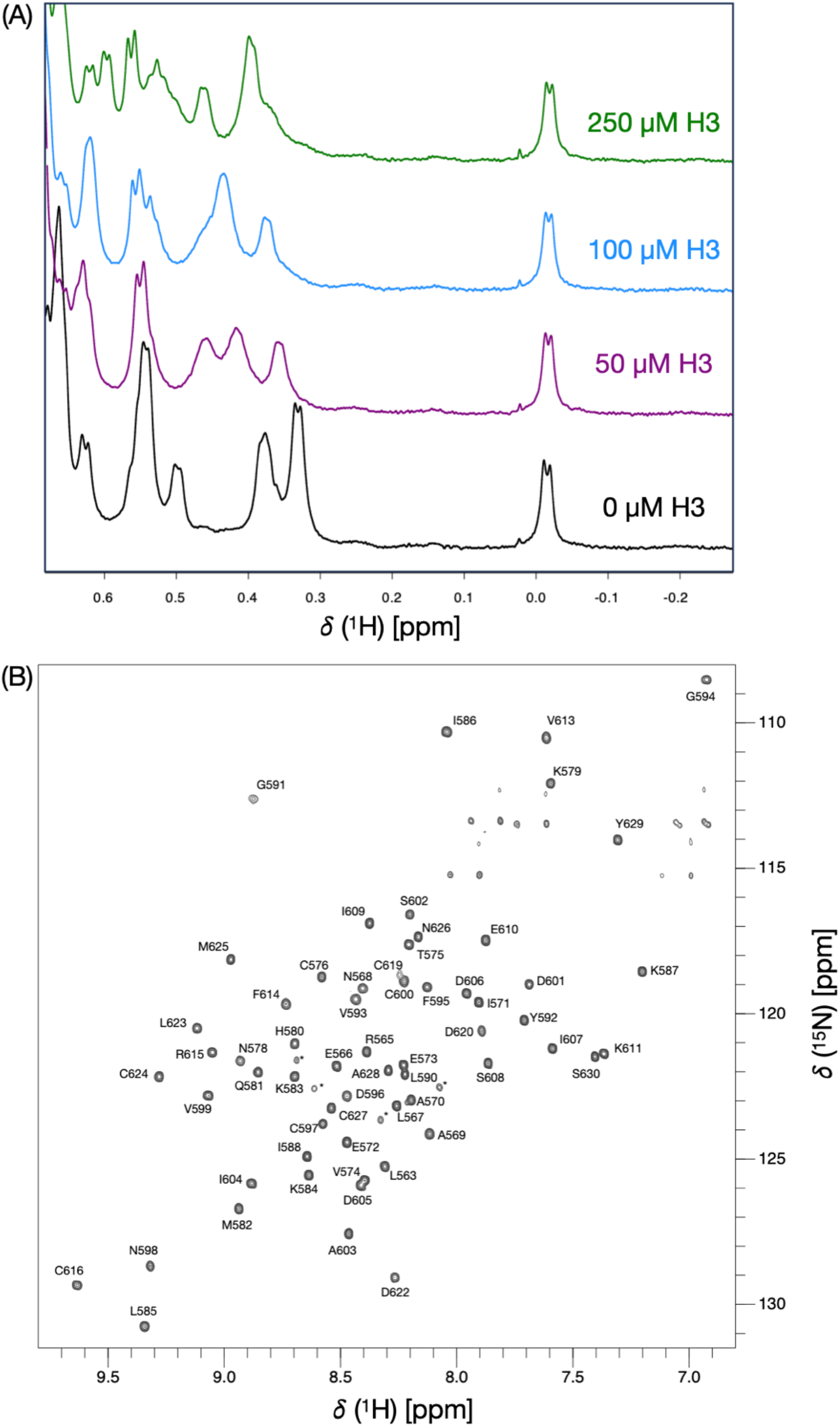
NMR spectra of KKT2 CL domain. (A) 750 MHz 1D ^1^H NMR spectra of the upfield methyl region of unlabeled KKT2 CL domain (KKT2^562–630^) showing changes in some methyl group peak positions upon the addition of histone H3^1–12^ peptide. These changes indicate an interaction between KKT2 CL and the H3 peptide. (B) 750 MHz ^1^H-^15^N BEST-TROSY spectrum of KKT2 CL in 25 mM HEPES, 100mM NaCl and 0.5 mM TCEP (95% H_2_O/5% D_2_O), at pH 7.2, 20°C. Peak assignments for backbone amides of 61 of the 66 non-proline residues are annotated. No amide peaks were identified for S562, H617, K618 and Y621. Four weaker peaks highlighted with * correspond to L563, R565, E566 and L567 in a minor species of KKT2 CL in which P564 adopts a cis conformation. Some peaks in the region of 110–114 ppm and upfield of ∼ 7.7 ppm are artefacts in the BEST-TROSY arising from incomplete cancellation of signal from the side chain amides of Asn and Gln. See also Figure S3A.

### NMR characterization of the structure and dynamics of KKT2 CL domain

To gain residue-specific information about this interaction, we obtained nearly complete resonance assignments for KKT2 CL using standard triple-resonance NMR protocols and isotope labeled KKT2 CL (Redfield, 2015). The 2D ^1^H-^15^N BEST-TROSY spectrum showed 65 well-dispersed peaks corresponding to backbone amides, consistent with a folded protein (Figure 3B). Resonances were assigned for 61 of the 66 non-proline residues; these assignments have been deposited in the BMRB (accession number 53855). Close examination of a 2D ^1^H-^15^N HSQC spectrum showed an additional backbone amide peak at 5.60 ppm/116.32 ppm which was assigned to I577 (Figure S3A). No amide peaks were identified for S562, H617, K618 and Y621. S562 is close to the N-terminus and may be absent due to rapid solvent exchange, while H617, K618 and Y621 may be broadened beyond detection due to slow conformational exchange.

Fast timescale (ps to ns) backbone dynamics of KKT2 CL were assessed using a {^1^H}-^15^N heteronuclear nuclear Overhauser effect (hetNOE) experiment (Kay et al., 1989). The majority of residues exhibit high {^1^H}-^15^N hetNOE ratios (>0.75), indicating a rigid backbone (Figure 4A). Increased flexibility is observed for N-terminal residues 563–574, C-terminal residues 629–630, and two loop regions (590–595 and 605–607). The random coil index order parameters (RCI S^2^) values, predicted by TALOS-N from the chemical shift data, agree well with the experimental hetNOE values (Figure 4A). Overall, most residues, including those within loop regions adopt a relatively rigid conformation on a fast timescale.

**Figure 4.**
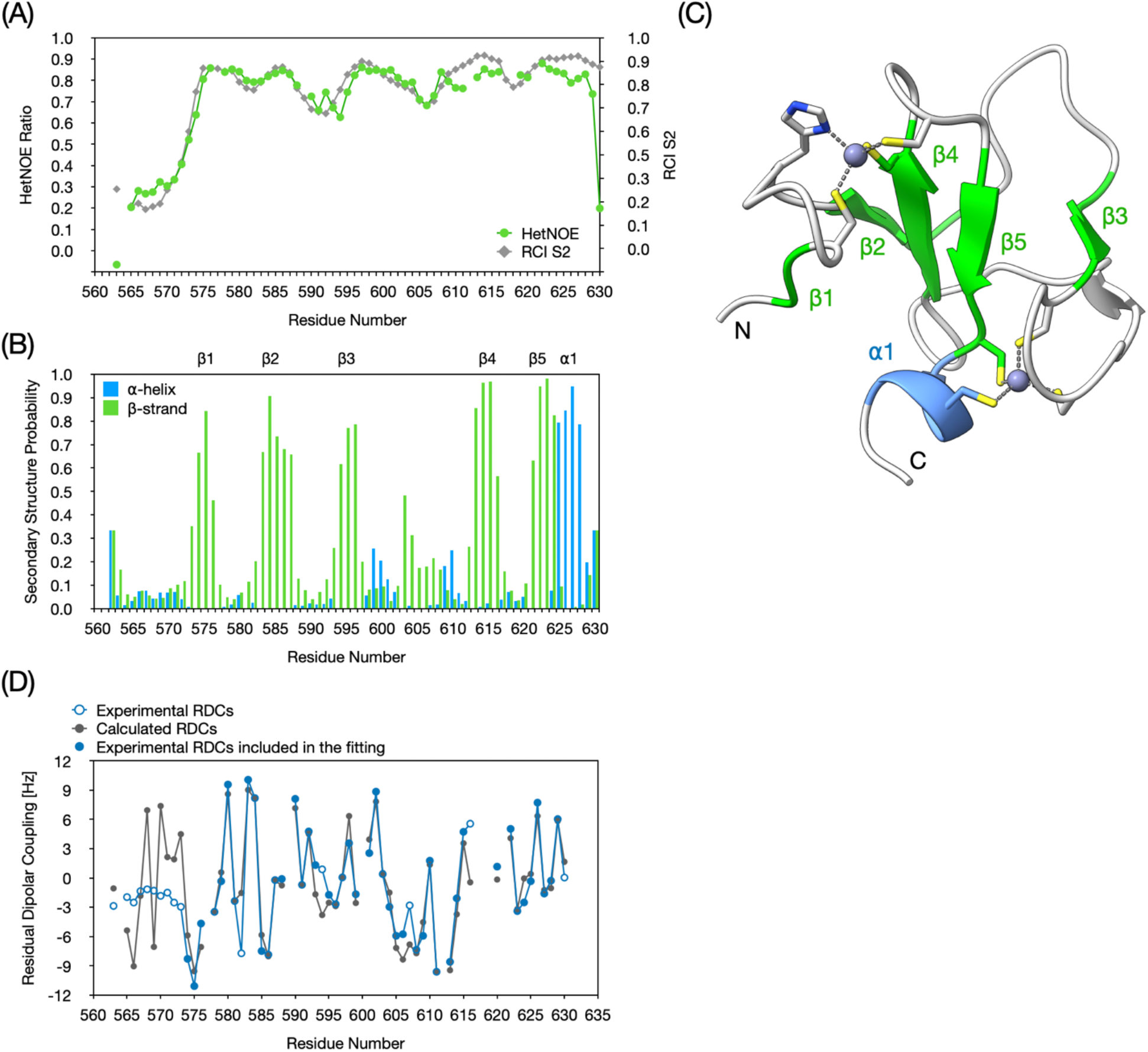
NMR characterization of KKT2 CL domain. (A) The {^1^H}-^15^N heteronuclear NOE ratios are plotted against the sequence of KKT2 CL. Most residues display hetNOE ratios >0.75, indicating a rigid conformation of the KKT2 CL domain backbone. The Random Coil Index order parameter (RCI S^2^) values obtained from TALOS-N analysis are also plotted. (B) TALOS-N secondary structure analysis of KKT2 CL. The probability of α-helix and b-strand secondary structure are shown in blue and green, respectively. A probability of >0.5 is usually used to assign a specific secondary structure type. TALOS-N predicts 5 β-strands and 1 α-helix in KKT2 CL. (C) The secondary structure elements identified using TALOS-N are highlighted in the cartoon representation of the AlphaFold3 model of KKT2 CL. The α-helix and β-strands are colored in blue and green, respectively. Zinc ions are shown as grey spheres and the side chains of the residues that coordinate the zinc ions are shown as sticks. (D) Experimental residual dipolar couplings (RDCs) (open and filled blue circles) are plotted as a function of the sequence of KKT2 CL. The small values for residues at the N-terminus are consistent with the flexibility of this region identified in the hetNOE experiment. RDC values used in the fitting procedure are shown as filled blue circles; RDCs calculated for all residues using the fitted alignment tensor are shown as grey filled circles and connected by a solid line. See also Figure S3.

Secondary structure elements were predicted from the assigned chemical shifts using TALOS-N (Shen and Bax, 2013). The analysis revealed five β-strands with an ⍺-helix at the C-terminus (Figure 4B). The core triple-stranded antiparallel β-sheet and the α-helix predicted by TALOS-N are in good agreement with the AlphaFold3 model of *T. brucei* KKT2 (Figure 4C). TALOS-N predicted β-strand structure for residues 594–596 but not for residues 601–604; both regions are predicted to form a short antiparallel β-sheet in the AlphaFold3 model (Figure 4C). Furthermore, TALOS-N predicted an additional β-strand involving residues 574–575 that was not predicted in the AlphaFold3 model (Figure 4C). TALOS-N identifies secondary structure on the basis of chemical shifts while visualization software, such as Chimera, uses a DSSP algorithm based on hydrogen bonds; this can lead to apparent discrepancies particularly for short regions of secondary structure. Analysis of through-space ^1^H-^1^H NOEs observed in NOESY spectra and the distances predicted from the AlphaFold3 model demonstrates that the solution structure of KKT2 CL is in good agreement with the AlphaFold3 model (Figure S3B and S3C).

Residual dipolar couplings (RDCs) are another NMR parameter that can be used to assess the quality of a structural model. RDCs for individual backbone amide groups depend on the orientation of the ^1^H_N_-^15^N bond vector with respect to an overall alignment tensor which describes the preferred orientation of the protein in the alignment medium. The experimentally measured RDCs can be compared with RDCs calculated from a structural model to assess the quality of the model. RDC values ranging from -11.1 Hz to +10.1 Hz were measured for KKT2 CL in 5% C12E6/hexanol (Figure 4D). Residues at the N-terminus of KKT2 CL have smaller experimental RDCs ranging from -3.0 to -1.2 Hz. This is consistent with the low {^1^H}-^15^N heteronuclear NOE values observed for these residues, which are typical of a disordered region of the polypeptide chain; these residues were excluded from further analysis. The molecular alignment tensor in the AlphaFold3 model was fitted to minimize the χ^2^ between the experimental and calculated RDCs for a set of 44 residues (see Materials and Methods for details); the Q value of 0.23 indicates good agreement between the experimental and predicted RDCs. The fitted alignment tensor was used to predict RDC values for all residues for which RDCs were measured (Figure 4D). Overall, this analysis of the residual dipolar couplings shows that the solution structure of KKT2 CL is in good agreement with the AlphaFold3 model; the level of agreement for the 44 residues that were fitted is consistent with the level of agreement often obtained when comparing RDCs measured in solution with those calculated from X-ray structures.

### NMR characterization of the interaction between KKT2 CL and histone H3 N-terminal peptide

To obtain residue-specific information about the KKT2 CL - histone H3 interaction, a series of 2D ^1^H-^15^N BEST-TROSY spectra of ^15^N-labelled KKT2 CL was collected upon addition of 50, 100 and 250 µM histone H3^1–12^ synthetic peptide. Overlay of the spectra reveals significant chemical shift changes for a subset of resonances (Figure 5A, S4A), indicating specific binding of the H3 peptide. Several resonances disappeared or broadened upon the initial addition of 50 µM H3 peptide, consistent with intermediate exchange on the NMR timescale. These residues likely undergo the largest chemical shift changes. To measure chemical shift changes for these residues, the titration experiment was repeated starting with a lower peptide concentration (5 µM) (Figure S5B); these spectra allow the initial direction of movement of peaks to be determined and allow the weak peaks that reappear at higher H3 concentration (250 µM) to be assigned to specific residues; to confirm these assignments a 3D ^15^N-edited NOESY-HSQC spectrum was collected with a high concentration of the H3 peptide. It is interesting to note that the peak corresponding to C619 becomes more intense as the peptide is added. In addition, at higher H3 peptide concentrations two new peaks appear in the BEST-TROSY spectra. These are assigned to K618 and Y621 which are broadened beyond detection in the absence of peptide (Figure S5B). Thus, binding of H3 peptide seems to remove the slow conformational exchange and stabilize the loop containing residues 618–621 which contains a Zn^2+^-coordinating residue C619. This region lies adjacent to the predicted H3-binding surface in the AlphaFold3 model and corresponds to the peptide-interacting loop in the p300 ZZ structure, consistent with a conserved binding mode.

**Figure 5.**
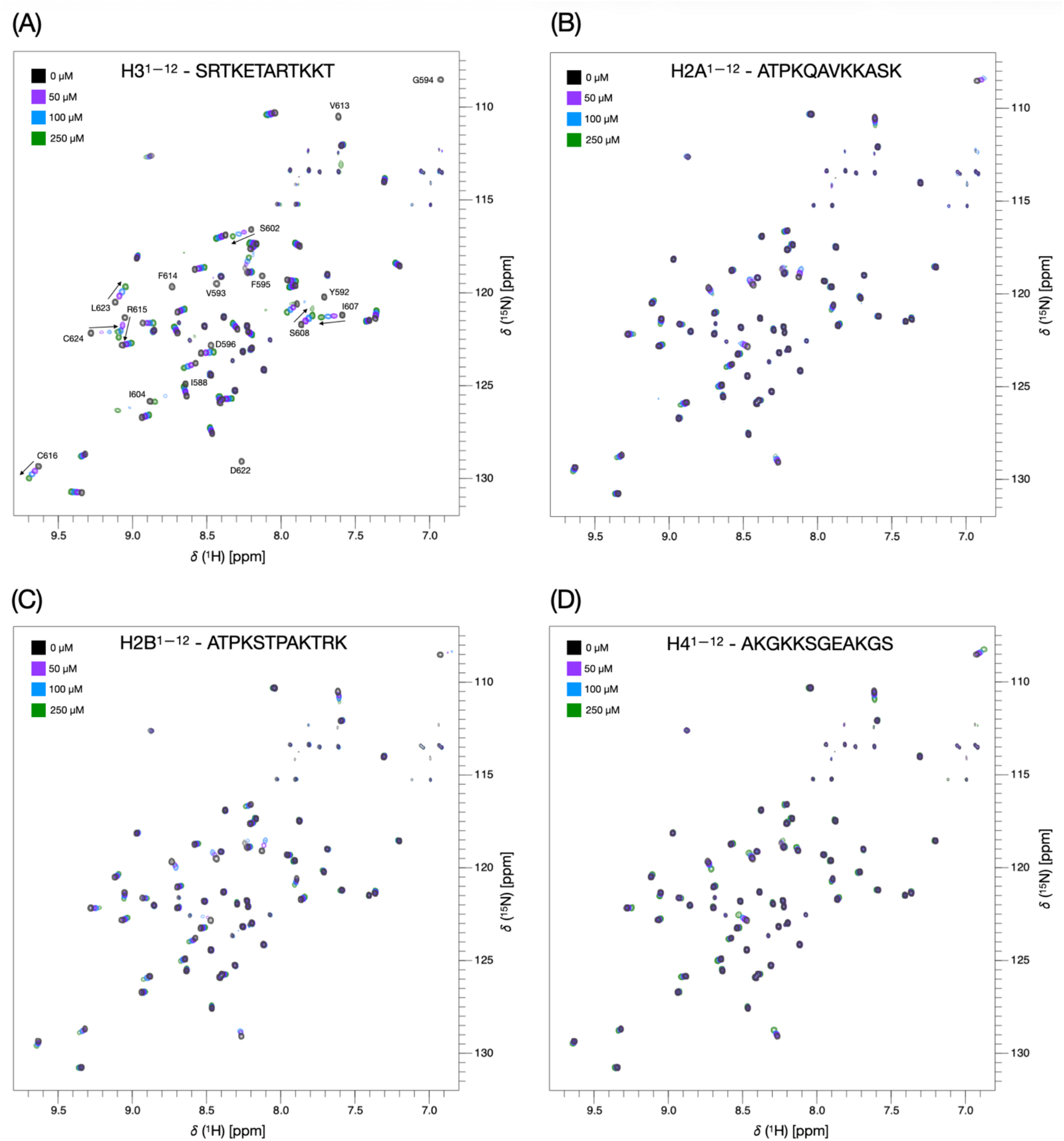
KKT2 CL domain binds histone H3 peptide. Overlay of 750 MHz ^1^H-^15^N BEST-TROSY spectra of 50 μM KKT2 CL domain (KKT2^562–630^ in 25 mM HEPES, 100mM NaCl and 0.5 mM TCEP at pH 7.2) in the absence (black) and with 50 (purple), 100 (blue) and 250 (green) μM of (A) H3^1–12^, (B) H2A^1–12^, (C) H2B^1–12^ and (D) H4^1–12^ N-terminal peptide added. For the H3 peptide in (A), peaks showing large chemical shift perturbations are labelled with their residue assignment and an arrow showing the direction of the peak movement. Direction of chemical shifts is not labelled for residues G594, D596 and D622, as their signals broaden and disappear upon the first addition of H3. See also Figure S5 and S6.

Similar results were obtained with shorter (H3^1–8^) and longer (H3^1–15^) peptides (Figure S5C,D). The observed chemical shift changes of these three versions of H3 are plotted against sequence in Figure S6. In contrast, much smaller chemical shift changes were detected upon addition of N-terminal peptides from histones H2A, H2B and H4 (Figure 5B–D, S4B–D), demonstrating much higher affinity and specificity of the KKT2 CL domain for histone H3. Among the affected residues, D622 exhibits one of the largest chemical shift perturbations; its peak disappears after addition of 10 µM H3 and only reappears as a very weak peak with 250 µM H3 (Figure S5B). Notably, D622 corresponds to the invariant aspartate that is critical for histone H3 binding in canonical ZZ domains (Mi et al., 2018). The invariant aspartate in H3-binding ZZ-domain binds the positively charged N-terminal amino-group of histone H3 (Mi et al., 2018). Consistent with this mechanism, residues showing the largest chemical shift changes cluster around D622 when mapped onto an AlphaFold3 model of the KKT2 CL-H3 complex, defining the predicted histone H3-binding pocket (Figure S7). The AlphaFold3 model is consistent with the NMR data and supports a binding mode in which the invariant aspartate coordinates the positively charged N-terminal amino group of histone H3. To directly test the role of D622 in H3 recognition, an isotope-labelled D622E mutant of KKT2 CL was generated and the NMR spectrum assigned (BMRB 53856). This mutation abolished the binding (Figure 6A,B), demonstrating that the invariant aspartate is essential for recognition of the histone H3 N-terminal tail. Together, these NMR data demonstrate that the KKT2 CL domain specifically binds the N-terminal tail of histone H3 *in vitro* and that this interaction depends on the invariant aspartate D622.

**Figure 6.**
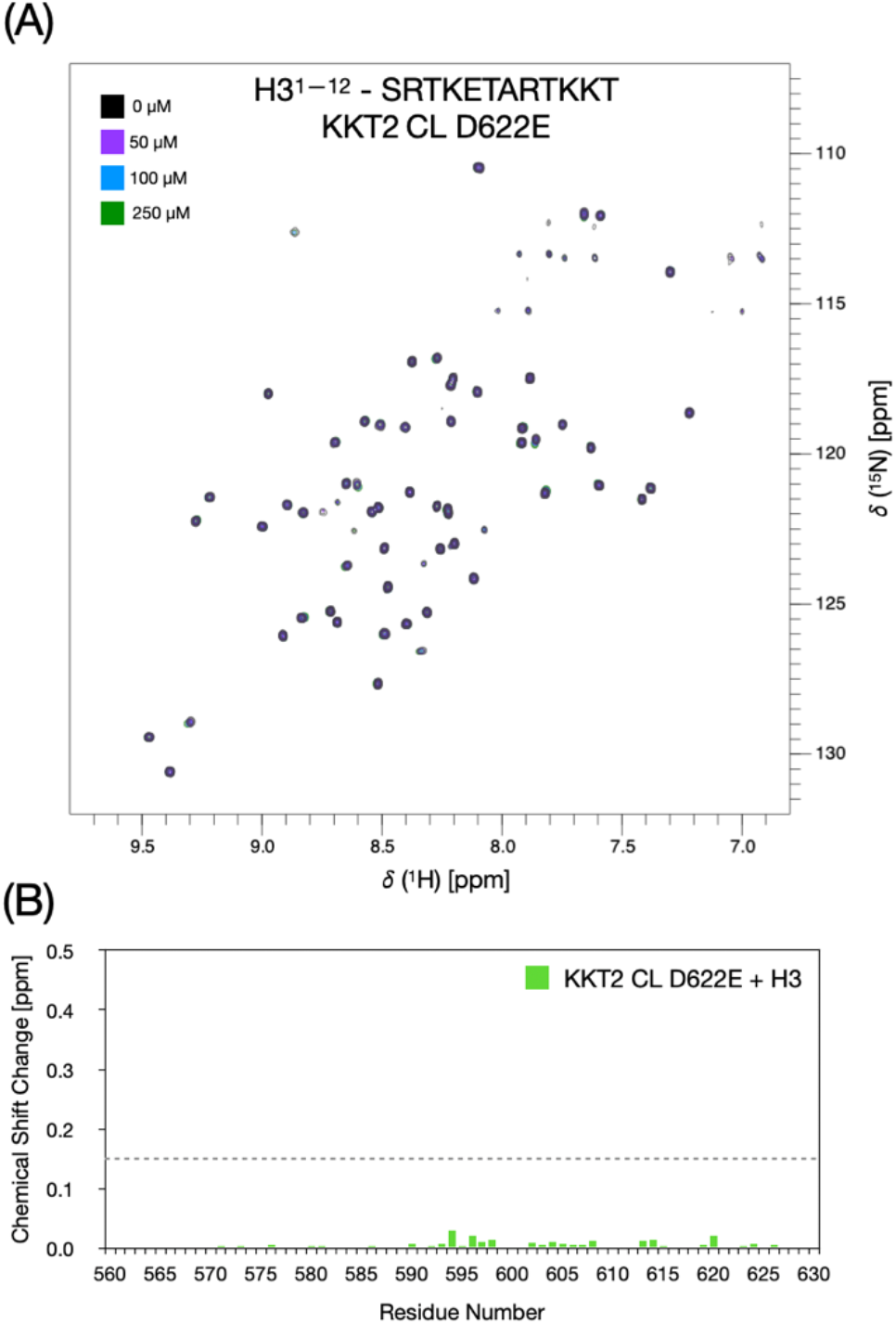
D622 is critical for the interaction of H3 with KKT2 CL. (A) Overlay of 750 MHz ^1^H-^15^N BEST-TROSY spectra of 50 μM KKT2 CL domain D622E (KKT2^562–630^ in 25 mM HEPES, 100mM NaCl and 0.5 mM TCEP at pH 7.2) in the absence (black) and with 50 (purple), 100 (blue) and 250 (green) μM of H3^1-12^ N-terminal peptide added. Comparison of the 2D ^1^H-^15^N BEST-TROSY spectra between wild-type and D622E mutant showed some changes in chemical shifts; these are likely to arise from relatively small changes in hydrogen bond lengths. Analysis of the ^1^Hα chemical shifts, which are sensitive to secondary structure, for wild-type and D622E KKT2 CL shows a mean difference of 0.03 ppm. The largest difference is observed for residue 622 but most of the difference of 0.34 ppm arises from the 0.32 ppm difference in expected Hα shift between aspartic and glutamic acids. Only residues G594, F595 and R615 have Hα shift differences of greater than 2 standard deviations from the mean (> 0.14 ppm); the sidechains of these residues pack against the sidechain of D622 and there may be small conformational adjustments needed to accommodate the longer sidechain of E622. The small Hα shift differences confirm that the D/E mutation does not alter the overall structure of KKT2 CL. (B) Quantification of chemical shift changes shown in (A) versus KKT2 sequence. Dotted line representing 0.15 ppm shows the cut-off of chemical shift changes for wild-type KKT2 CL that were mapped on the structure shown in Figure S7.

### PTMs on histone H3 side chains fail to enhance the interaction

Because histone H3 is not specific to centromeres, the identified interaction between KKT2 CL and histone H3 would not explain how the CL domain specifically localizes at centromeres. We therefore hypothesized that the interaction between the CL domain and histone H3 could be regulated by post-translational modifications (PTMs) on the histone tail. We tested nine different PTMs on the side chains of highly conserved residues: phosphorylation of T3 and T6, mono/di-methylation^symmetric^/di-methylation^asymmetric^ of R2, and mono/di/tri-methylation or acetylation of K4. However, none of these modifications increased the affinity (Figure 7, S8). T3 phosphorylation weakens binding of H3 to KKT2 CL, indicating that introduction of a negative charge at this position destabilizes the protein-peptide complex (Figure 7E, S8E).

**Figure 7.**
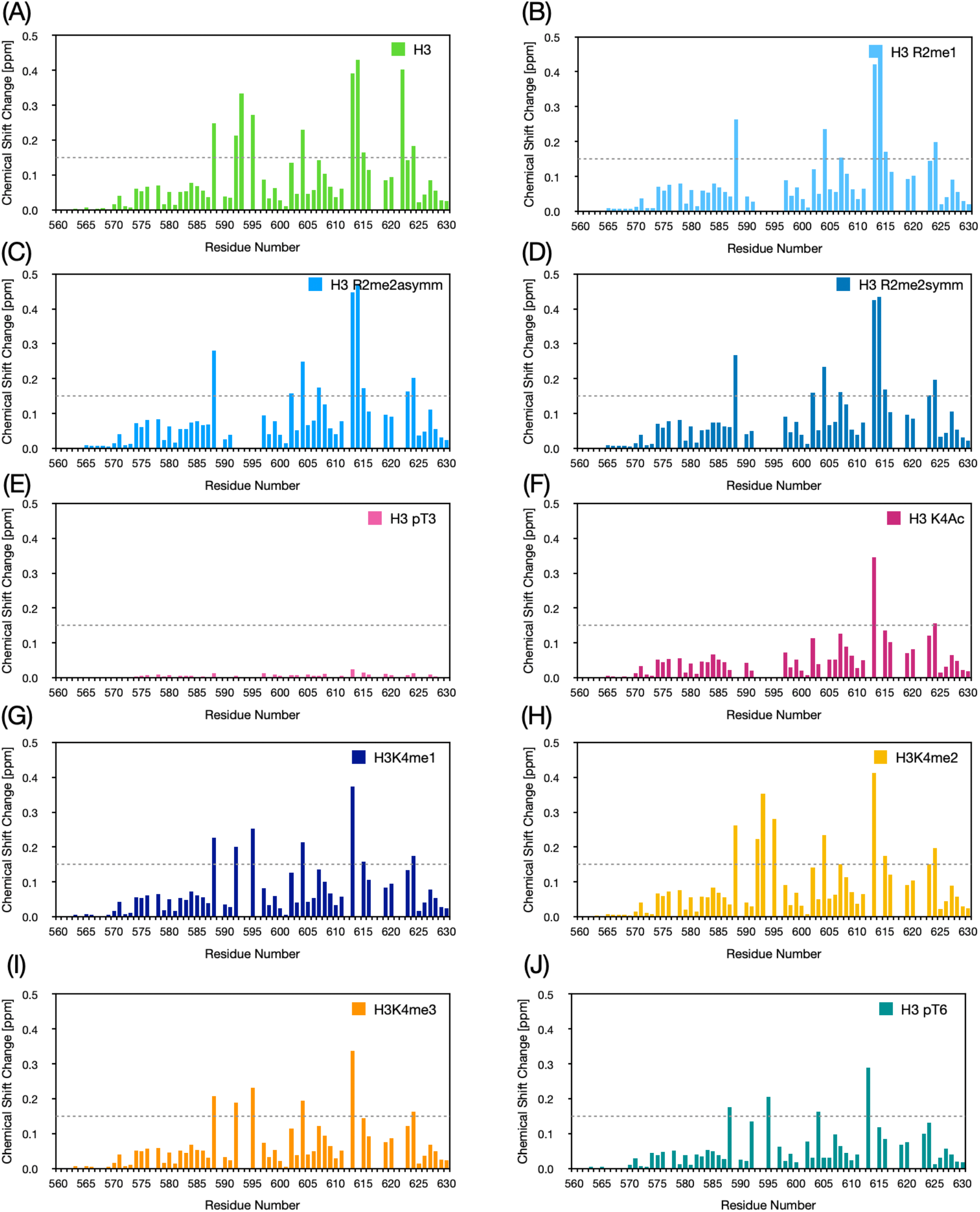
Effect of PTMs on H3 binding. Quantification of chemical shift changes upon addition of 250 μM H3 peptide versus KKT2 sequence for modified histone H3 peptides. (A) Unmodified H3, (B) R2 monomethylation, (C) R2 asymmetric dimethylation, (D) R2 symmetric dimethylation, (E) T3 phosphorylation, (F) K4 acetylation, (G) K4 monomethylation, (H) K4 dimethylation, (I) K4 trimethylation, (J) T6 phosphorylation. Dotted line representing 0.15 ppm shows the cut-off of chemical shift changes for unmodified H3 that were mapped on the structure shown in Figure S7. For all plots, no chemical shift change is shown for I577, P589, P612, H617, K618 or Y621 due to the absence of peaks for these residues in the BEST-TROSY spectrum collected in the absence of peptide. In some plots, no chemical shift change is shown for Y592, V593, G594, F595, D596, I604 and D622 because the peaks corresponding to these residues are still broadened beyond detection in the presence of 250 μM peptide indicating an interaction. See also Figure S8.

### Interaction between KKT2 CL and histone H3 is abolished by Nα modifications

Besides side chain modifications, we also tested two modifications at the Nα, namely acetylation and methylation. The N-terminal acetylation is a common PTM in eukaryotic proteins, which neutralizes the positive charge of the NH_3_ ^+^ group (Aksnes et al., 2016). Nα acetylation of the interacting peptides has been shown to inhibit the interaction with ZZ domains (Mi et al., 2018; Zhang et al., 2018a). This is consistent with the notion that ZZ domains use the negatively-charged aspartate to recognize the positively-charged NH_3_ ^+^ group of the target (Zhang et al., 2018a; Mi et al., 2018; Liu et al., 2020; Tencer et al., 2022). Similarly, AlphaFold3 predicted interaction between the aspartate of KKT2 CL and the positively-charged Nα of the Ser1 residue (Figure S7). In contrast to the unmodified histone H3 peptide, no major chemical shift changes were observed upon addition of N-terminally acetylated histone H3 to ^15^N-KKT2 CL, indicating a lack of interaction (Figure 8A,B). We next tested Nα methylation of histone H3, which is an abundant modification in *T. brucei* (Kraus et al., 2020). Strikingly, mono-methylation of histone H3 N-terminus also resulted in no major chemical shift changes indicating a lack of interaction (Figure 8C,D). Because methylation has little effect on the charge of the NH_3_^+^ group, we hypothesize that the modified N-terminus causes a steric clash preventing H3 docking onto the CL domain.

**Figure 8.**
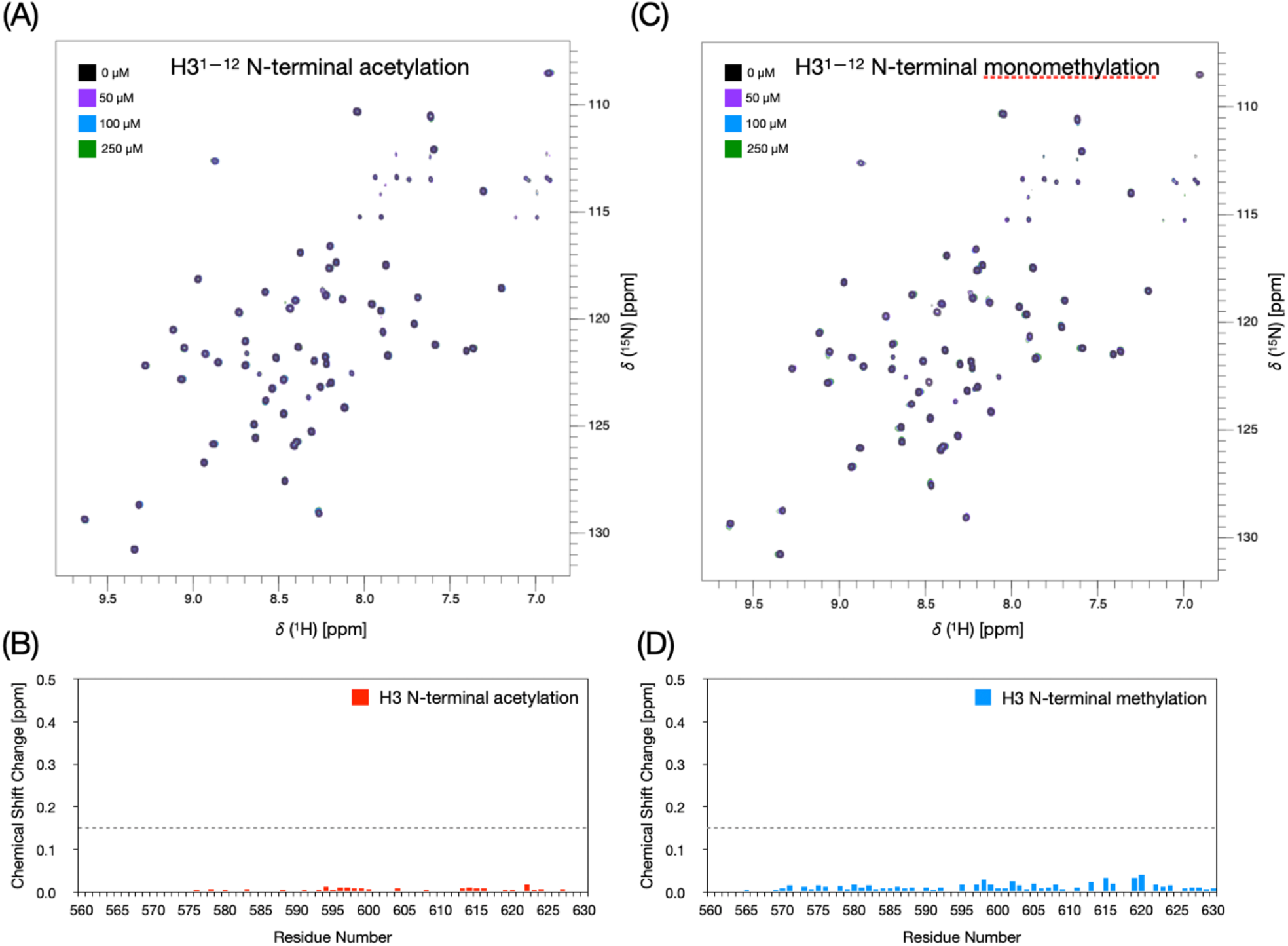
Nα acetylation and methylation of histone H3 abolish interaction with KKT2 CL. (A) Overlay of 750 MHz ^1^H-^15^N BEST-TROSY spectra of 50 μM KKT2 CL domain (KKT2^562–630^ in 25 mM HEPES, 100mM NaCl and 0.5 mM TCEP at pH 7.2) in the absence (black) and with 50 (purple), 100 (blue) and 250 (green) μM of H3^1-12^ peptide with N-terminal acetylation added. (B) Quantification of chemical shift changes shown in (A) versus KKT2 sequence. Dotted line representing 0.15 ppm shows the cut-off of chemical shift changes for unmodified H3 that were mapped on the structure shown in Figure S7. (C) Overlay of 750 MHz ^1^H-^15^N BEST-TROSY spectra of 50 μM KKT2 CL domain (KKT2^562–630^ in 25 mM HEPES, 100mM NaCl and 0.5 mM TCEP at pH 7.2) in the absence (black) and with 50 (purple), 100 (blue) and 250 (green) μM of H3^1-12^ peptide with N-terminal monomethylation added. (D) Quantification of chemical shift changes shown in (C) versus KKT2 sequence. Dotted line representing 0.15 ppm shows the cut-off of chemical shift changes for unmodified H3 that were mapped on the structure shown in Figure S7.

We next used isothermal titration calorimetry (ITC) to assess thermodynamic parameters of the interaction between KKT2 CL and unmodified H3 peptide. KKT2 CL showed moderate affinity for H3^1–12^ with a dissociation constant (K_d_) in the micromolar range (K_d_ = 59.9 ± 3.7 µM; Figure 9A). The ITC result revealed a 1:1 stoichiometry for the complex. The interaction was exothermic and predominantly enthalpy-driven (ΔH = −5.60 ± 0.69 kcal/mol), although entropy also contributed favorably (-TΔS= −0.17 kcal/mol. Derived ΔS= 0.55 cal/mol·K). In contrast, no interaction was observed between unmodified H3^1–12^ and the KKT2 CL D622E variant (Figure 9B) nor between Nα-monomethylated H3^1–12^ and wild-type KKT2 CL (Figure 9C), consistent with the NMR analysis (Figure 6A,B, 8C,D). Taken together, these results show that the interaction between KKT2 CL and histone H3 is highly sensitive to any changes in the interaction interface.

**Figure 9.**
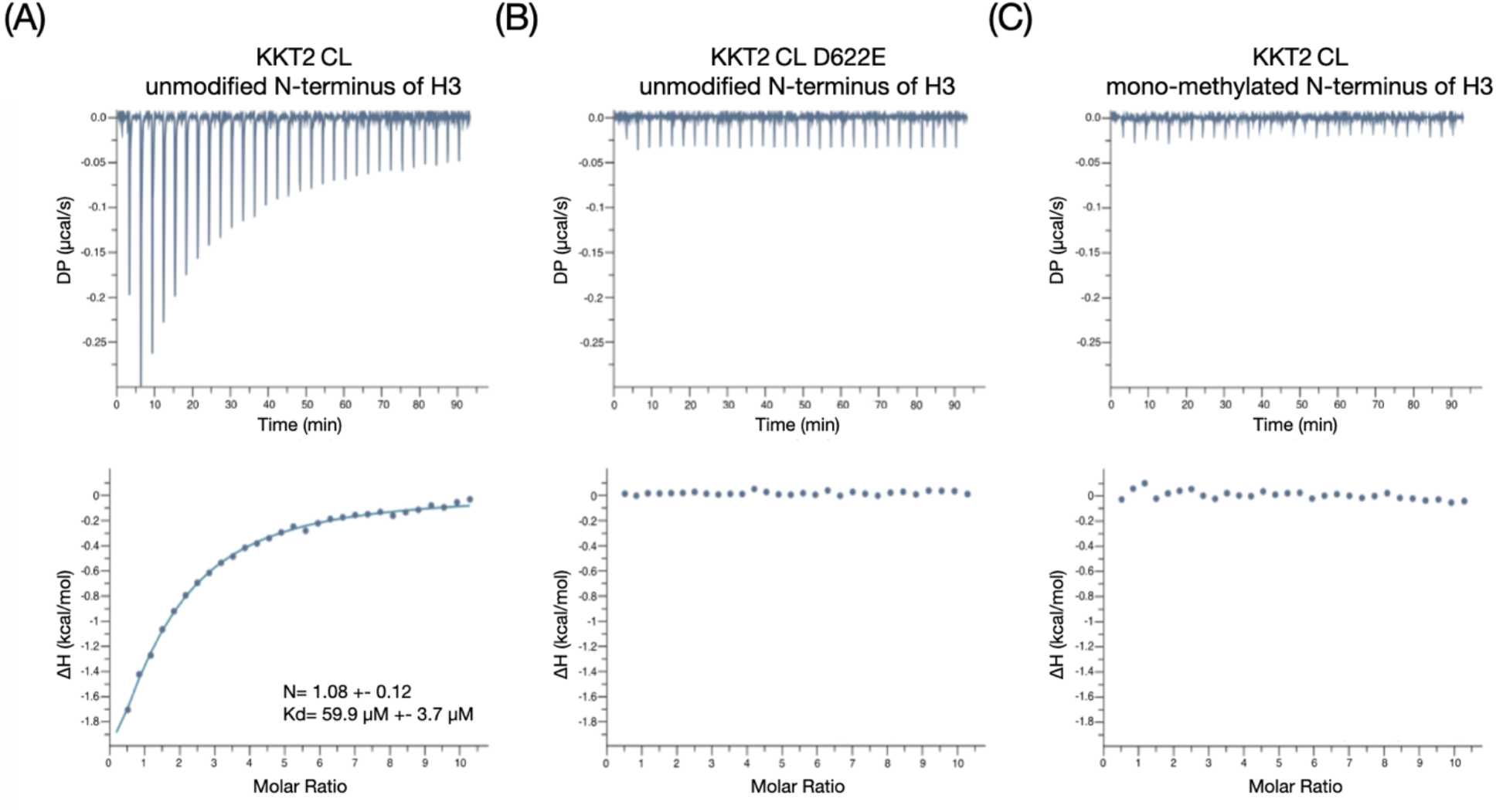
ITC analysis of the interaction between KKT2 CL and the N-terminus of histone H3. (A) KKT2 CL interacts with unmodified histone H3^1–12^. (B) KKT2 CL D622E mutant does not interact with unmodified histone H3^1–12^. (C) KKT2 CL does not interact with Nα mono-methylated histone H3^1–12^. Top panels show the raw ITC data; bottom panels show the integrated heat data corrected for heat of dilution and fit to a standard 1:1 binding model (Malvern Instruments MicroCal Origin software, v1.3). DP, differential power.

## Discussion

It has long been known that kinetoplastids lack CENP-A (Lowell and Cross, 2004; Berriman et al., 2005), leaving the mechanism of kinetochore specification enigmatic. Identification of their kinetochore proteins did not solve this question (Akiyoshi and Gull, 2014). The findings reported in this manuscript finally shed light on this long-standing question in trypanosome biology. Our main findings are that 1) KKT2 CL has structural similarity to ZZ, 2) KKT2 CL binds the N-terminus of histone H3, and 3) the interaction is abolished by Nα modifications of histone H3. Although we cannot exclude the possibility that untested PTMs (or combinations of PTMs) increase the affinity, none of the side-chain PTMs tested in this study resulted in increased affinities. We did find that phosphorylation of T3, but not of T6, reduces the interaction; this PTM has not been observed for H3 in trypanosome cells so this result might not be biologically relevant. However, this result may indicate that the hydroxyl group of T3 interacts with KKT2 CL.

It has been known for two decades that, in *T. brucei*, the majority of histone H3 molecules are modified at its Nα. Edman sequencing of histones isolated from *T. brucei* failed to provide information for histone H3 (Janzen et al., 2006). Consistent with this, mass spectrometry analysis showed that Nα of histone H3 is heavily methylated (Kraus et al., 2020). Together with our finding that even mono-methylation of the histone H3 N-terminus abolishes the binding to KKT2 CL, we propose a hypothesis that kinetochore specification relies on the absence of Nα modifications of histone H3 specifically at the centromere. In other words, binding of KKT2 CL to non-centromeric regions is prevented by Nα methylations and acetylation of histone H3. It will be crucial to test this possibility by determining the modification status of centromeric histone H3 in trypanosome cells.

The CL domain is also present in KKT3. Although the biochemical analyses presented here were limited to KKT2 because recombinant KKT3 could not be obtained in a form suitable for structural or biophysical characterization, our previous analyses suggest that the mechanism described here is likely conserved. In particular, KKT3 has an invariant aspartate (D692 in *T. brucei*), whose mutation abolished the centromere localization of KKT3 (Marcianò et al., 2021). It is therefore conceivable that the KKT3 CL domain binds the N-terminus of histone H3, although this possibility needs to be tested through biochemical investigation.

It will also be important to identify the methyltransferase(s) responsible for histone H3 Nα methylations. Compared to Nα acetylation that occurs on many eukaryotic proteins, much less is known about Nα methylations (Aksnes et al., 2016; Demetriadou et al., 2020). Thus far, only two types of Nα methyltransferases are known in eukaryotes, METTL11A/B (also known as NRMT1/2 or NTMT1/2) and METTL13, both of which have a strict substrate specificity (Wong and Eirin-Lopez, 2021; Diaz et al., 2021; Chen et al., 2023). METTL11A/B target >100 substrates that have a X_aa_-Pro-Lys/Arg consensus motif at the N-terminus (Tooley et al., 2010; Webb et al., 2010). For example, the N-terminal glycine of human CENP-A (Gly-Pro-Arg) is tri-methylated by METTL11A, which is important for robust recruitment of kinetochore proteins to centromeres (Bailey et al., 2013; Sathyan et al., 2017). By contrast, METTL13 has only one known target, eEF1a (Gly-Lys-Glu-Lys) (Jakobsson et al., 2018). *T. brucei* histone H3 (Ser-Arg-Thr-Lys) does not match either of these consensus motifs, hinting at the presence of a distinct type of methyltransferase. Further studies are needed to identify the responsible methyltransferase and dissect the mechanism of kinetochore specification in kinetoplastids.

## Materials and Methods

### Expression and purification of KKT2^562–630^ and KKT2^562–630^ D622E for NMR

The *T. brucei* KKT2 CL^562–630^ (pBA1631) construct was amplified from a KKT2-containing plasmid using oligonucleotides BA1563 and BA2113, and cloned into the RSF_Duet-1 vector (Novagen) with the NEBuilder HiFi DNA Assembly Cloning Kit (New England Biolabs). This construct was originally designed for crystallization trials, and the glutamic acid residue at position 630 was substituted with serine. The KKT2 CL^562–630^ D622E mutant (pBA3056) was generated from pBA1631 using oligonucleotides BA2556 and BA2557 via a site-directed mutagenesis reaction.

*E. coli* BL21 (DE3) cells were transformed with the respective plasmids and grown overnight in 2xTY medium. The following day, 50 mL of the overnight culture was harvested, washed with 25 mL of M9 minimal medium, and resuspended in 30 mL of M9 medium. From this suspension, 5 mL was inoculated into 500 mL of M9 minimal medium in a 2.5 L flask, supplemented with 0.1 mM ZnSO_4_ and 1 g/L ^15^NH_4_Cl. At an optical density (OD_600_) of approximately 0.6, the temperature was lowered to 20 °C, and IPTG was added 30 min later for overnight induction. For double labelling of KKT2^562–630^, the M9 medium was additionally supplemented with 4 g/L ^13^C-glucose. A total of 6 L of culture medium was used for each preparation.

Protein extraction and purification were performed as previously described (Marcianò et al., 2021). Briefly, cells were pelleted and resuspended in lysis buffer (25 mM HEPES pH 7.5, 150 mM NaCl, 1 mM tris(2-carboxyethyl)phosphine [TCEP], 10 mM imidazole, and 1.2 mM PMSF) and lysed using a French press. After centrifugation (48,000 × g for 30 min at 4 °C), the clarified lysate was applied to TALON affinity resin, washed with lysis buffer, and eluted by gravity flow with elution buffer (25 mM HEPES pH 7.5, 150 mM NaCl, 1 mM TCEP, and 250 mM imidazole). The eluate was incubated overnight at 4 °C with TEV protease for tag cleavage (as a result, Gly-Ser from the tag precedes the KKT2 sequence). The protein was further purified by anion-exchange chromatography (Resource Q or HiTrap Q FF, Cytiva) using a 0.05–1 M NaCl gradient. Fractions containing the protein of interest were pooled, concentrated, and subjected to size-exclusion chromatography (HiPrep Superdex 75 16/60, GE Healthcare) in 25 mM HEPES pH 7.5, 100 mM NaCl, and 1 mM TCEP. The purified protein was concentrated and stored at −80 °C. Final concentrations were 522 µM for ^15^N-labeled KKT2^562-630^, 260 µM for ^15^N-labeled KKT2^562-630^ D622E, and 90 µM for ^15^N,^13^C-labeled KKT2^562-630^. The presence of zinc ions in purified proteins was confirmed using zinc assay (Merck, MAK032).

### Expression and purification of KKT2^562–630^ and KKT2^562–630^ D622E for ITC

*E. coli* BL21 (DE3) cells were transformed with pBA1631 or pBA3056 and grown overnight in 2xTY medium. The following day, 10 mL of the overnight culture was inoculated into 1 L 2xTY medium. At an optical density (OD_600_) of 0.6 – 0.8, the temperature was lowered to 20 °C, and IPTG was added for overnight induction.

For protein purification, cells were pelleted and resuspended in lysis buffer (25 mM HEPES pH 7.5, 500 mM NaCl, 0.5 mM TCEP, 5 mM imidazole) supplemented with 500 U/L benzonase nuclease, 20 µg/mL leupeptin, 20 µg/mL pepstatin, 20 µg/mL E-64, and 0.4 mM PMSF, and lysed by sonication. After centrifugation (48,000 × g for 30 min at 4 °C), the clarified lysate was applied to TALON affinity resin, washed with lysis buffer, and eluted by gravity flow with elution buffer (25 mM HEPES pH 7.5, 150 mM NaCl, 250 mM imidazole, 0.5 mM TCEP). The eluate was incubated overnight at 4 °C with TEV protease for His-tag cleavage and dialyzed into 25 mM HEPES, 150 mM NaCl, 5 mM imidazole, and 0.5 mM TCEP. The protein was further purified by anion-exchange chromatography (HiTrap Q, Cytiva) using a 0.05–1 M NaCl gradient. Fractions containing the protein of interest were pooled, concentrated, and subjected to size-exclusion chromatography (Superdex 75 Increase 10/300 GL, Cytiva) in 25 mM HEPES pH 7.5, 150 mM NaCl, and 0.5 mM TCEP. The purified protein was concentrated and stored at −80 °C.

#### NMR spectroscopy and analysis of NMR data

All NMR samples were prepared in 25 mM HEPES pH 7.2, 100 mM NaCl, 0.5 mM TCEP, 95% H_2_O/5% D_2_O. All NMR spectra were acquired at 20 °C using either 750 or 950 MHz spectrometers equipped with Bruker Avance III HD consoles and 5 mm TCI CryoProbes. Resonance assignments for KKT2^562–630^ were obtained using 2D ^1^H-^13^C HSQC, ^1^H-^15^N HSQC and ^1^H-^15^N BEST-TROSY (Schulte-Herbrüggen and Sorensen, 2000; Lescop et al., 2007) experiments and 3D experiments including ^15^N-edited NOESY-HSQC, ^15^N-edited TOCSY-HSQC, ^13^C-edited NOESY-HSQC, (H)CC(CO)NH, HC(C)H-TOCSY and BEST-TROSY versions of HNCA, HNCO, HN(CO)CACB, HNCACB, and HN(CA)CO. The HNCACB, and ^13^C-edited NOESY-HSQC were collected at 950 MHz; all other 3D data were collected at 750 MHz. Backbone ^1^H^N^-^15^N and ^1^H^α^ resonance assignments for the D622E variant of KKT2^562–630^ were obtained using ^1^H-^15^N HSQC, ^1^H-^15^N BEST-TROSY (Schulte-Herbrüggen and Sorensen, 2000; Lescop et al., 2007) experiments and a 3D ^15^N-edited NOESY-HSQC.

All 3D NMR data, except the ^15^N-edited NOESY-HSQC and TOCSY-HSQC for WT KKT2 CL, were collected with 25% non-uniform sampling in the two indirect dimensions using standard Bruker sampling schedules. 2D NMR data were processed using NMRPipe (Delaglio et al., 1995) and 3D NUS data were processed with the hmsIST software (Hyberts et al., 2012) and NMRPipe. Spectra were analysed and assignments recorded using CcpNmr Analysis version 2.5 (Vranken et al., 2005). ^1^H and ^13^C chemical shifts were referenced using DSS and ^15^N chemical shifts were referenced indirectly. Details of the specific experiments and sample conditions can be found in the BMRB deposition files (53855 and 53856 for WT and D622E KKT2 CL, respectively). ^1^H, ^13^C and ^15^N chemical shifts of KKT2^562–630^ were analysed using TALOS-N (Shen and Bax, 2013) to predict secondary structure propensities.

The {^1^H}-^15^N heteronuclear NOE was measured for a 0.5 mM sample of KKT2^562–630^ using the standard HSQC-based heteronuclear NOE experiment recorded with and without ^1^H saturation for 3 sec at 750 MHz (Kay et al., 1989). The {^1^H}-^15^N NOE was calculated as the ratio of the peak intensities in the spectra recorded with and without ^1^H saturation. Peak heights were determined using CCPN Analysis (Vranken et al., 2005).

Residual dipolar couplings were measured for partially aligned KKT2^562–630^. Isotropic ^1^H^N^-^15^N splittings were measured for a 100 mM sample of ^15^N-labeled KKT2^562–630^. Partial alignment of KKT2^562–630^ was achieved using C12E6/*n*-hexanol liquid crystals prepared as described previously (Rückert and Otting, 2000). Briefly, a 15% C12E6/*n-* hexanol stock solution was prepared in HEPES buffer (25 mM HEPES, 100 mM NaCl, pH 7.5). 100 μl of this stock solution was added to 200 μl of the 100 μM ^15^N-labelled KKT2^562–630^ to achieve a final concentration of 5% C12E6/*n-*hexanol. ^1^H^N^-^15^N splittings for both samples were measured using BEST-TROSY and semi-BEST-TROSY experiments at 20℃ (Schulte-Herbruggen and Sorensen 2000, Lescop, Schanda et al. 2007). RDCs were measured as the difference between the splitting observed in the isotropic and aligned data sets. Three measurements were taken in each experiment and the average RDC value was calculated.

RDC values were measured for 59 of the 61 assigned non-proline residues in KKT2 CL. The principal components and orientation of the molecular alignment tensor were fitted to minimize the χ^2^ between the experimental and calculated RDCs using the KKT2 CL AlphaFold3 model to which ^1^H had been added using X-PLOR version 3.8 (Brünger, 1992). Eleven residues with {^1^H}-^15^N heteronuclear NOE values of less than 0.6, indicative of backbone flexibility, were excluded from the fitting procedure; this includes residues 563, 565-573 at the N-terminus and 630 at the C-terminus. This resulted in a data set of 48 experimental RDC values. Q values were calculated to assess the quality of the fits between experimental and calculated RDCs using the method of Cornilescu and co-workers (Cornilescu et al., 1998). A Q value of 0.35 was obtained from this initial analysis; four residues, M582, G594, I607 and C616, contribute significantly to this Q value with differences between experimental and calculated RDCs ranging from 4.2 to 5.2 Hz. If these four residues are removed from the data set then the 44 remaining RDCs give a Q value of 0.23 with D_a_ and R values of 5.7 and 0.53, respectively.

#### NMR data collection with histone titrations

Peptides used in this study are listed in Table S1. Histone peptides synthesized by Designer Bioscience or Thermo Fisher were dissolved in 25 mM HEPES pH 7.5, 100 mM NaCl and 0.5 mM TCEP to prepare 10 mM stock solutions. For NMR experiments, 350 µL samples containing 50 µM ^15^N-KKT2^562–630^ were prepared in 25 mM HEPES pH 7.5, 100 mM NaCl, 0.5 mM TCEP, 5% (v/v) D_2_O. Histone peptides were titrated into the protein sample to achieve final peptide concentrations of 50, 100 and 250 µM. In addition, lower concentrations of 5, 10, 20 and 40 µM were collected for the unmodified H3 peptide. The pH was measured and adjusted after each addition of peptide to avoid pH-dependant changes in the collected spectra. Samples were incubated for approximately 10 min at room temperature prior to acquisition of a 2D ^1^H-^15^N BEST-TROSY spectrum. Combined chemical shift changes observed between peak positions in the absence of and in the presence of 250 μM peptide were calculated in CcpNmr Analysis using ((Δ^1^H^N^)^2^ + (0.15 x Δ^15^N)^2^)^1/2^. Residues with shift changes of > 0.15 ppm are indicated by the dashed line in the plots. A 3D ^15^N-edited NOESY-HSQC spectrum was collected for KKT2 CL in the presence of 1 mM H3 peptide; this allowed assignments for peaks that disappeared and then reappeared during the titration to be confirmed.

#### N-terminal tail peptides of histone H3 for ITC

Lyophilized histone H3^1–12^ peptides with or without Nα mono-methylation were synthesized by Peptide Synthetics and dissolved in 25 mM HEPES pH 7.5, 150 mM NaCl and 0.5 mM TCEP to prepare 10 mM stock solution. For ITC experiments, the peptide concentration was measured on Nanodrop using Abs214. Peptides were dialyzed into 25 mM HEPES pH 7.5, 150 mM NaCl and 0.5 mM TCEP using Pur-A-Lyzer Midi 1000 (Sigma-Aldrich) at 4 °C overnight, together with proteins KKT2^562–630^ and KKT2^562–630^ D622E. Final concentration of 2 mM peptides and 30 µM proteins were used for ITC.

#### Isothermal titration calorimetry (ITC)

ITC experiments were performed using MicroCal Auto-iTC200 at 25 °C. Samples had been dialyzed into 25 mM HEPES pH 7.5, 150 mM NaCl, 0.5 mM TCEP at 4 °C overnight. 120 µL of 2 mM unmodified or mono-methylated H3^1–12^ was injected into 400 µL of 30 µM KKT2^562–630^ or KKT2^562–630^ D622E in a total of 30 injections (1 x 0.5 μL and 29 x 1.0 μL), at intervals of 180 sec, with stirring at 750 rpm. Heat of dilution was measured by titrating peptide into buffer and subtracted from raw thermograms during analysis. Data were integrated and fitted into a one-site binding model using MicroCal PEAQ-ITC and corrected for heat of dilution to obtain apparent binding stoichiometry (N), dissociation constant (Kd), enthalpy (ΔH), and entropy (−TΔS). Then, −TΔS was used to calculate ΔS. Protein and peptide concentrations were measured on Nanodrop using absorbance at 280 and 214 nm respectively.

### SEC-MALS

Size-exclusion chromatography (ÄKTA-PURE 25 M; Cytiva) coupled with UV, static light scattering and refractive index (RI) detection (Viscotec SEC-MALS 20 and Viscotek RI Detector VE3580; Malvern Instruments, Malvern, Worcestershire, UK) were used to determine the absolute molecular mass of KKT2^562–630^ in solution. Injections of 100 μL of 8 mg/mL of the protein were run on a calibrated Superdex 75 Increase 10/300 GL (Cytiva) size exclusion column pre-equilibrated in 25 mM HEPES pH 7.5, 150 mM NaCl, 0.5 mM TCEP at 22 °C with a flow rate of 1.0 mL/min. Light scattering, RI and A280 nm were analysed by a homo-polymer model (Omnisec, v5.02; Malvern Instruments) using the following parameters for KKT2^562–630^: ∂A280 nm/∂c = 0.56 AU·mL−1·mg−1, ∂n/∂c = 0.190 mL·g−1 and buffer RI value of 1.336.

#### *In silico* structure and interaction predictions

Structural predictions of *T. brucei* KKT2 CL, KKT2 CL with H3, and KKT3 CL were performed with AlphaFold3 (Abramson et al., 2024). All structure figures were made using ChimeraX (Meng et al., 2023).

#### Multiple sequence alignment

Protein sequences of histone H3 were retrieved from UniProt (UniProt Consortium, 2019), TriTryp database (Aslett et al., 2010), or a previous publication (Butenko et al., 2020). Multiple sequence alignment was performed with MAFFT (L-INS-i, version 7) (Katoh et al., 2019) and visualized with the Clustalx coloring scheme in Jalview (version 2.11) (Waterhouse et al., 2009).

## Acknowledgments

We thank Edinburgh Protein Production Facility (EPPF) for helping with ITC experiments. Bungo Akiyoshi was supported by a Wellcome Discovery Award (227243/Z/23/Z). Christopher W Wood was funded by a BBSRC sLOLA grant (BB/X003027/1). The Department of Biochemistry NMR Facility has benefitted from funding provided by the Edward Penley Abraham Fund, the John Fell Fund and the Wellcome Trust.

## Author contributions

A.C., P.L., G.M., W.A., and M.I. purified recombinant proteins. P.L., G.M., and C.R. performed and analyzed NMR experiments. A.C. performed ITC experiments. P.L., G.M., S.F., C.W.W., C.R. and B.A. performed in silico structural predictions and analyses. A.C., P.L., G.M., C.R., and B.A. designed experiments and wrote the manuscript with input from all authors.

## Rights retention

This research was funded in whole, or in part, by the Wellcome Trust (227243/Z/23/Z). For the purpose of open access, the author has applied a CC BY public copyright licence to any Author Accepted Manuscript version arising from this submission.

## Supporting figures and figure legends

**Figure S1.**
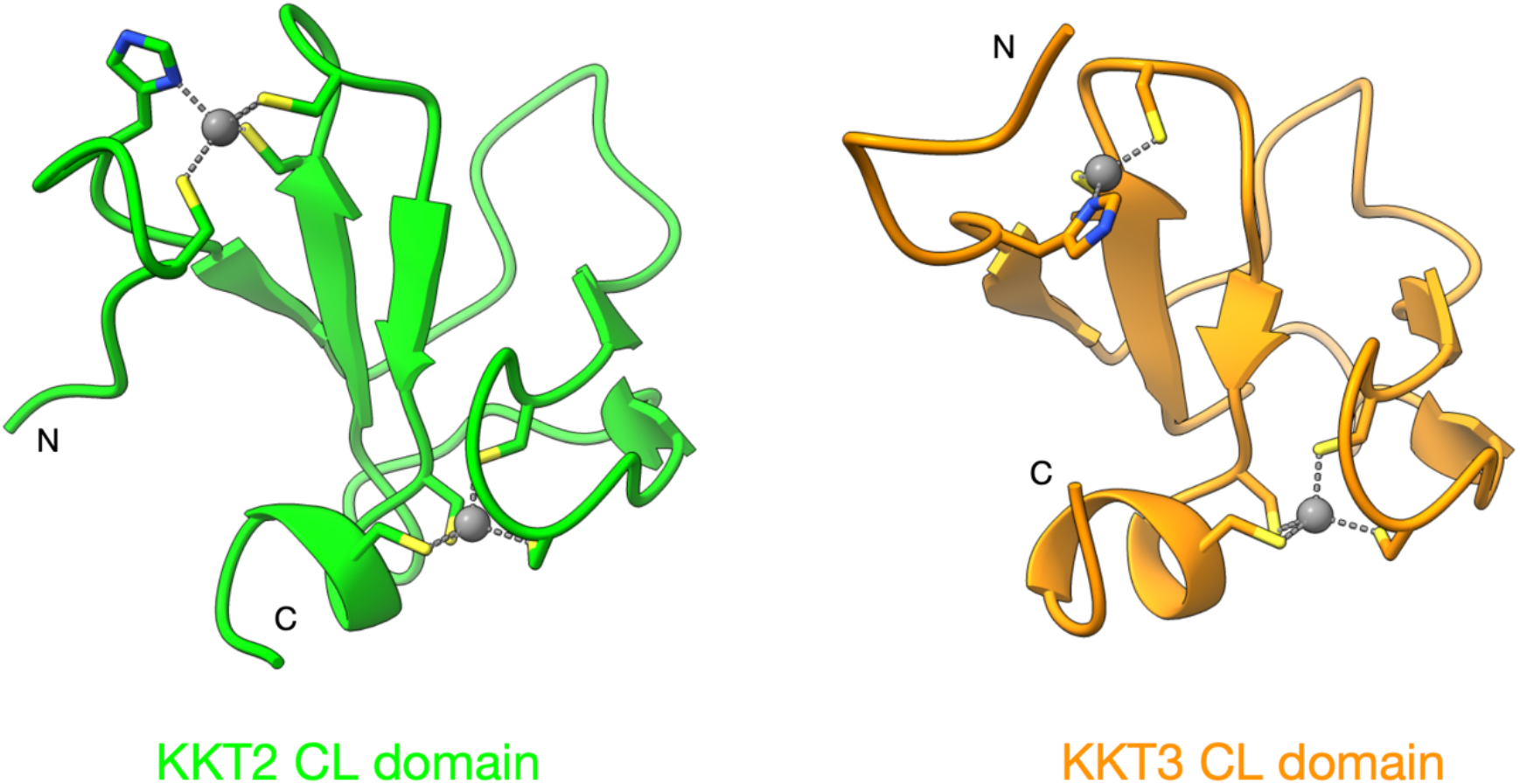
AlphaFold3 models of KKT2 CL and KKT3 CL domains show similar folds. Structures of *T. brucei* KKT2 CL^562–630^ (green) and KKT3 CL^645–701^ (orange) were predicted using AlphaFold3 in the presence of two zinc ions. Residues that coordinate zinc ions (grey) are shown as sticks and colored by heteroatom. KKT2 residues 562–570 are not shown for clarity.

**Figure S2.**
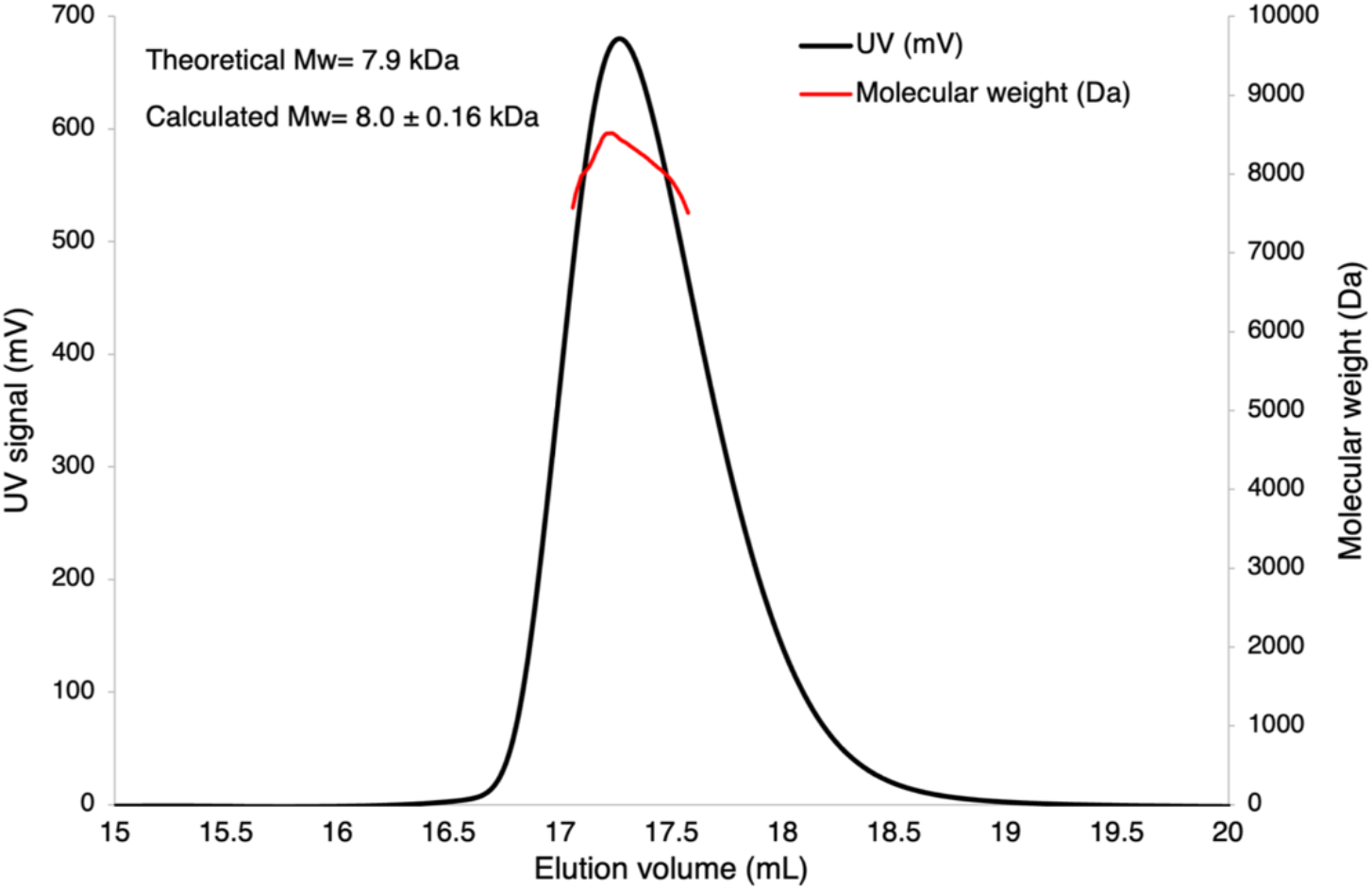
KKT2 CL is monomeric. SEC-MALS analysis of KKT2 CL. The measured molecular mass (8.0 ± 0.16 kDa) closely matched the theoretical monomer mass (7.9 kDa), indicating that KKT2 CL is monomeric in solution.

**Figure S3.**
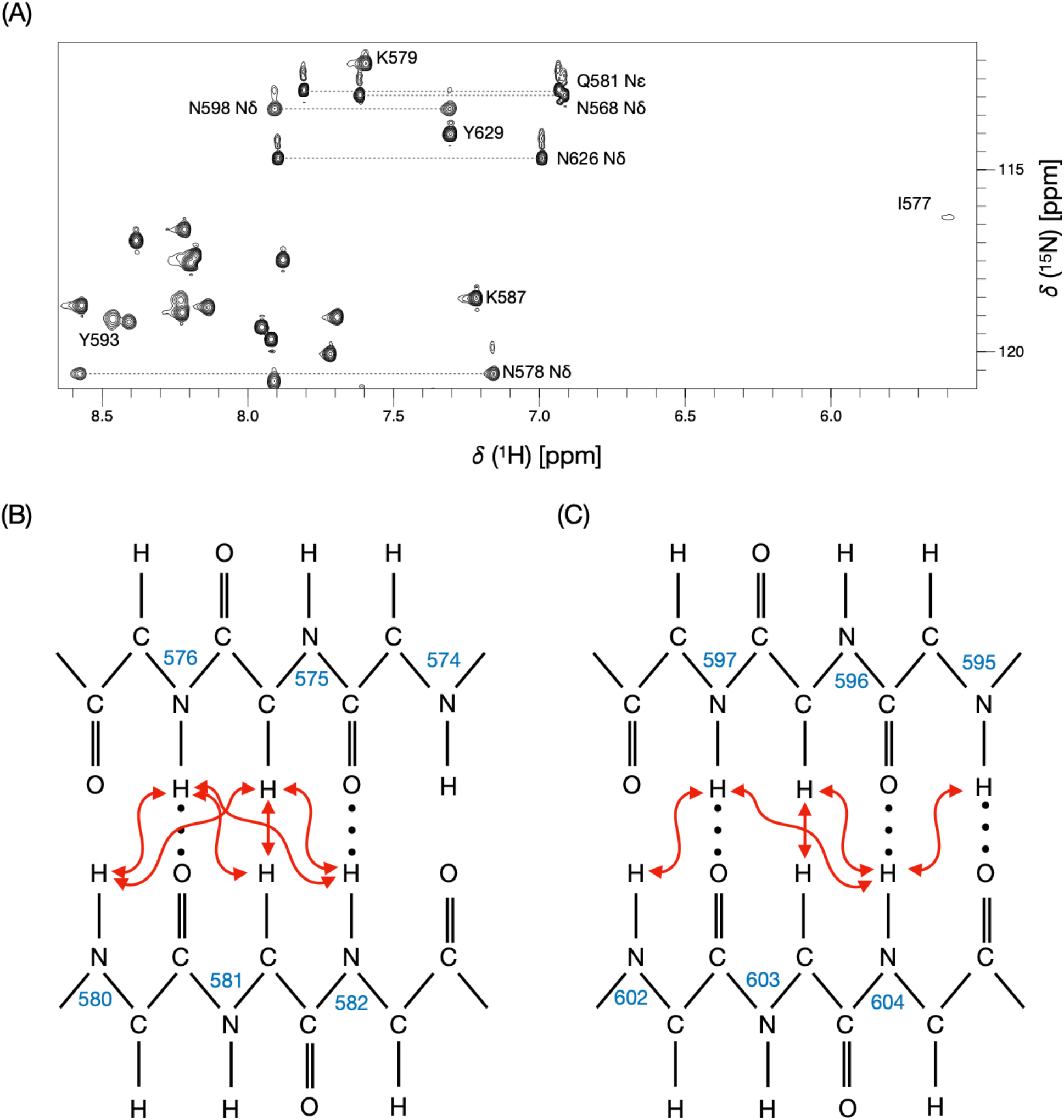
^1^H-^15^N HSQC spectrum of KKT2 CL and long-range NOEs observed for KKT2 CL. (A) Expansion of a region from the 750 MHz ^1^H-^15^N HSQC spectrum of KKT2 CL in 25 mM HEPES, 100mM NaCl and 0.5 mM TCEP (95% H_2_O/5% D_2_O), at pH 7.2, 20 °C. The pairs of peaks corresponding to the side chain NH_2_ groups of the three asparagine and one glutamine residues are labelled. In addition, the upfield shifted backbone amide peak of I577 at 5.6 ppm is labelled. This peak is not observed in the BEST-TROSY spectrum due to the shaped ^1^H pulses used in this experiment. (B) Schematic representation of the b-sheet interactions observed for residues 574–576 and 580–582. TALOS-N predicted a β-strand involving residues V574–T575 that was not predicted in the AlphaFold3 model. Analysis of the latter structure shows hydrogen bonds between M582 N and V574 O and between C576 N and H580 O (illustrated with the dotted lines); these are consistent with an antiparallel interaction between residues 574–576 and 580–582 in the AlphaFold3 model. These are probably not assigned as b-strands because the DSSP algorithm would also expect V574 N – M582 O and H580 N – C576 O hydrogen bonds in a regular antiparallel b-sheet, which are not observed, and the f and y torsion angles deviate from ideal values. TALOS-N does not predict a b -strand for residues 580–582. It is interesting to note that the side chains of C576 and H580 coordinate a Zn^2+^ ion and this may lead to a distortion of the structure that affects chemical shifts and the TALOS-N prediction of secondary structure. The 3D ^15^N-edited and ^13^C-edited NOESY spectra show ^1^H-^1^H NOEs involving the backbone H^N^ and Ha of residues 575, 576 and residues 580, 581, 582 consistent with an antiparallel b-sheet (shown as red arrows); the T575 Ha to Q581 Ha NOE is particularly characteristic of an anti-parallel b-sheet structure. These NOEs are also predicted from the AlphaFold3 structure. (C) Schematic representation of the b-sheet interactions observed for residues 595–597 and 602–604. TALOS-N predicted β-strand structure for residues 594–596 but did not predict β-strand structure for residues 601–604; both these regions are predicted as b-strands involved in an antiparallel interaction in the AlphaFold3 model. Four hydrogen bonds are observed between F595 N and I604 O, I604 N and F595 O, C597 N and S602 O, D601 N and C597 O illustrated with the dotted lines); the first pair are consistent with regular antiparallel b-sheet while the second pair indicate a more distorted structure. It is interesting to note that the side chains of C597 and C600, which are located in the turn connecting the two b-strands, coordinate a Zn^2+^ ion. The 3D NOESY spectra show ^1^H-^1^H NOEs involving the backbone H^N^ and Ha of residues 595, 596, 597 and residues 602, 603, 604 consistent with an antiparallel b-sheet (shown as red arrows); the D596 Ha to A603 Ha NOE is particularly characteristic of an anti-parallel b-sheet structure. These NOEs are also predicted from the AlphaFold3 structure.

**Figure S4.**
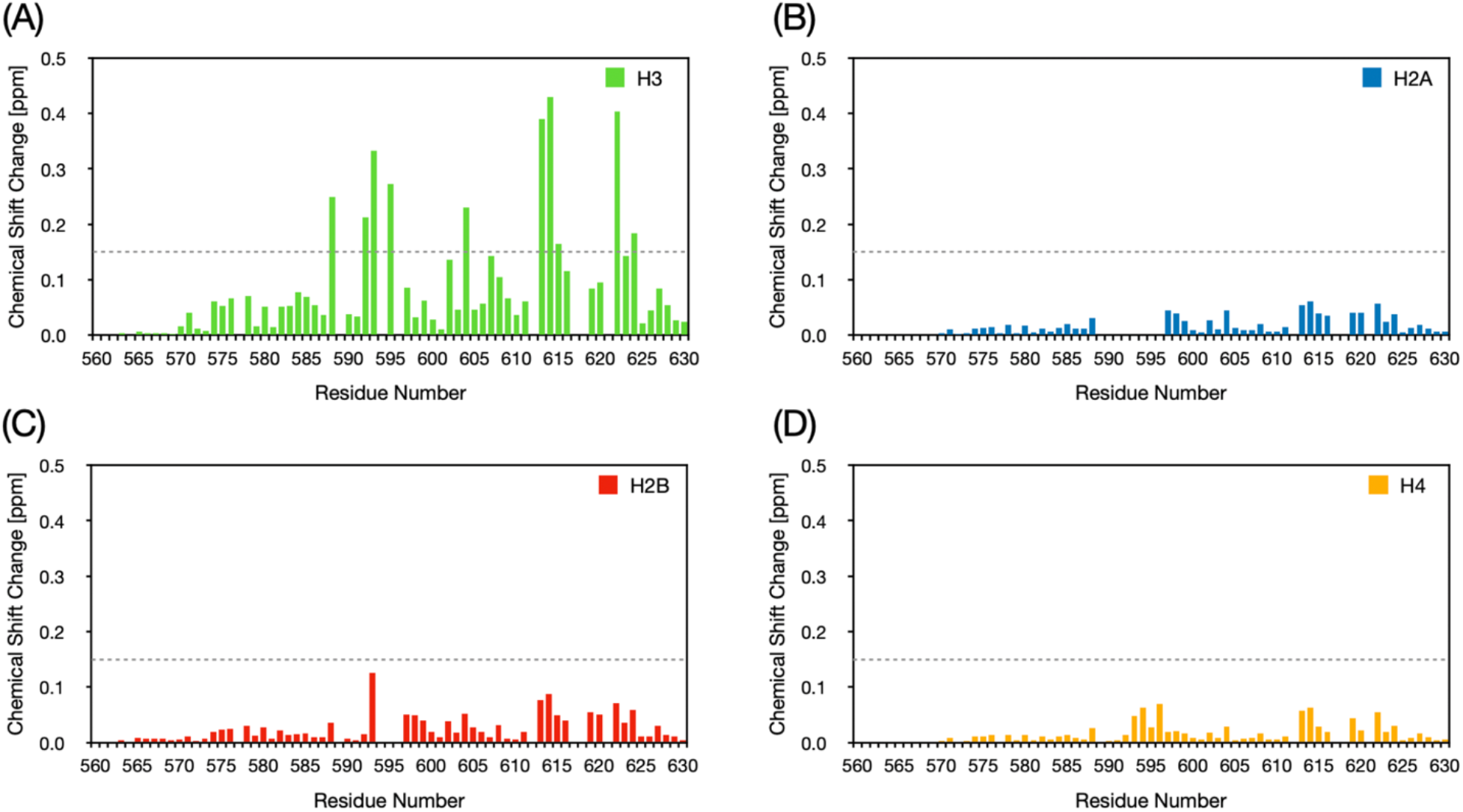
KKT2 CL domain binds histone H3 peptide. Bar plots show the magnitude of chemical shift changes (ppm) as a function of KKT2 sequence for histone peptides (A) H3, (B) H2A, (C) H2B and (D) H4. Note that Figure S4A is the same as Figure 7A, and is shown here for comparison. The dashed horizontal line at 0.15 ppm indicates the threshold used to identify significantly perturbed residues. Residues exceeding this cut-off were mapped onto the KKT2 AlphaFold3 model (Figure S7). For all plots, no chemical shift change is shown for I577, P589, P612, H617, K618 or Y621 due to the absence of peaks for these residues in the BEST-TROSY spectrum collected in the absence of peptide. In some plots, no chemical shift change is shown for residues 592–596, 604 and 622 because the peaks corresponding to these residues are still broadened beyond detection in the presence of 250 μM peptide. See also Figure S5 and S6.

**Figure S5.**
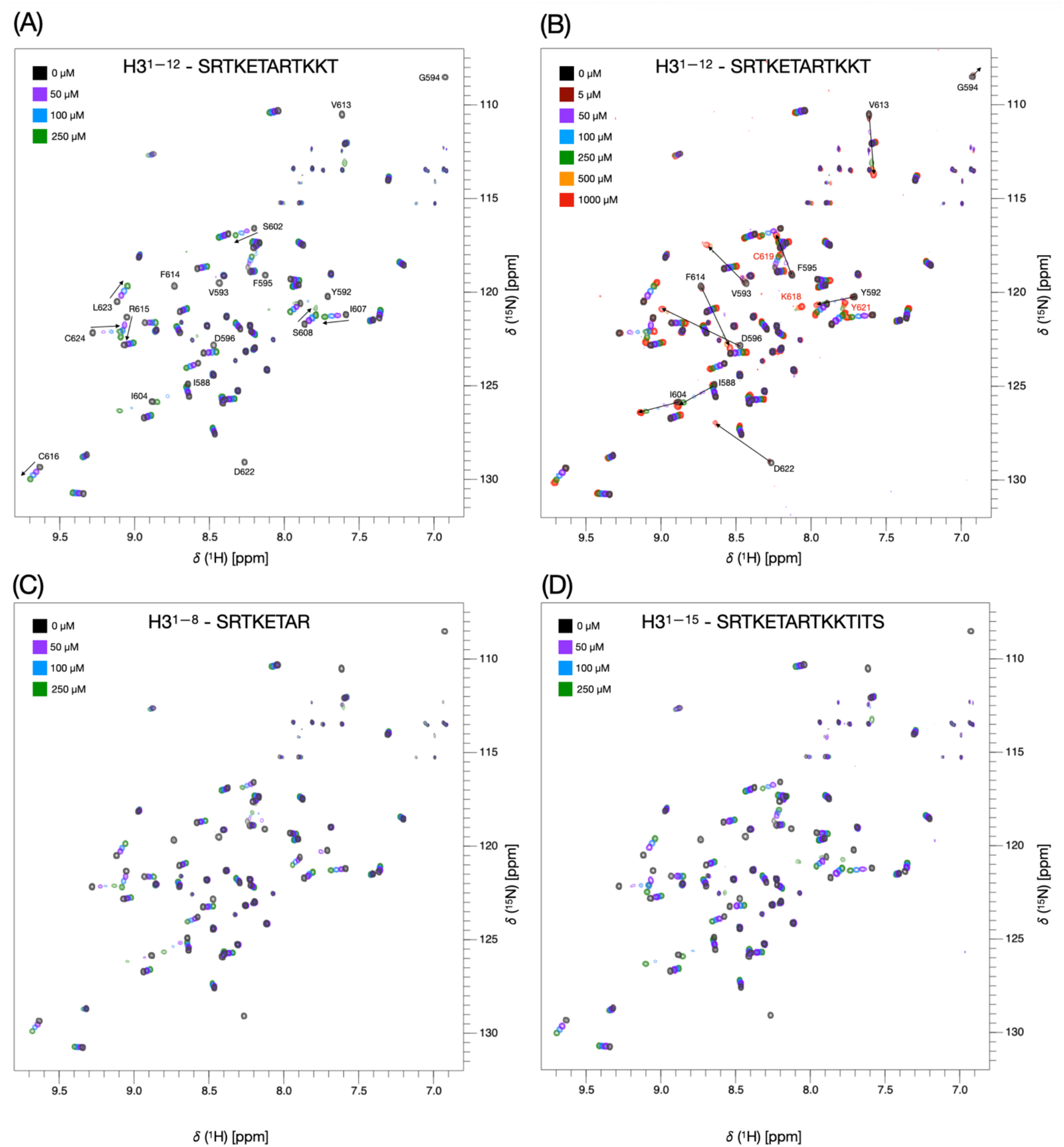
NMR titration of KKT2 CL with different lengths of histone H3 peptide. Overlay of 750 MHz ^1^H-^15^N BEST-TROSY spectra of 50 μM KKT2 CL domain (KKT2^562–630^ in 25 mM HEPES, 100mM NaCl and 0.5 mM TCEP at pH 7.2) in the absence (black) and with 50 (purple), 100 (blue) and 250 (green) μM of histone H3 N-terminal peptide containing (A/B) 12, (C) 8 and (D) 15 residues added. Panel (B) also shows spectra collected with 5μM (maroon), 500μM (orange) and 1mM (red) peptide. In (A), peaks showing large chemical shift perturbations are labelled with their residue assignment and an arrow showing the direction of the peak movement. Note that Figure S5A is the same as Figure 5A, and is shown here for comparison. In (B), many peaks which disappear in (A) following addition of 50 μM H3 can be seem in the 5 μM spectrum allowing the direction in which the peaks shift to be identified. Peaks for these residues reappear and become stronger in the presence of 500 μM and 1 mM H3 peptide when the KKT2 CL binding site is fully saturated. The peak corresponding to C619, labelled in red, becomes stronger at higher concentrations of H3 peptide. Two new peaks which are visible in the spectra collected with 500μM and 1 mM peptide can be assigned to K618 and Y621. These residues are located in the same Zn^2+^ binding loop as C619; binding of H3 appears to stabilize this loop.

**Figure S6.**
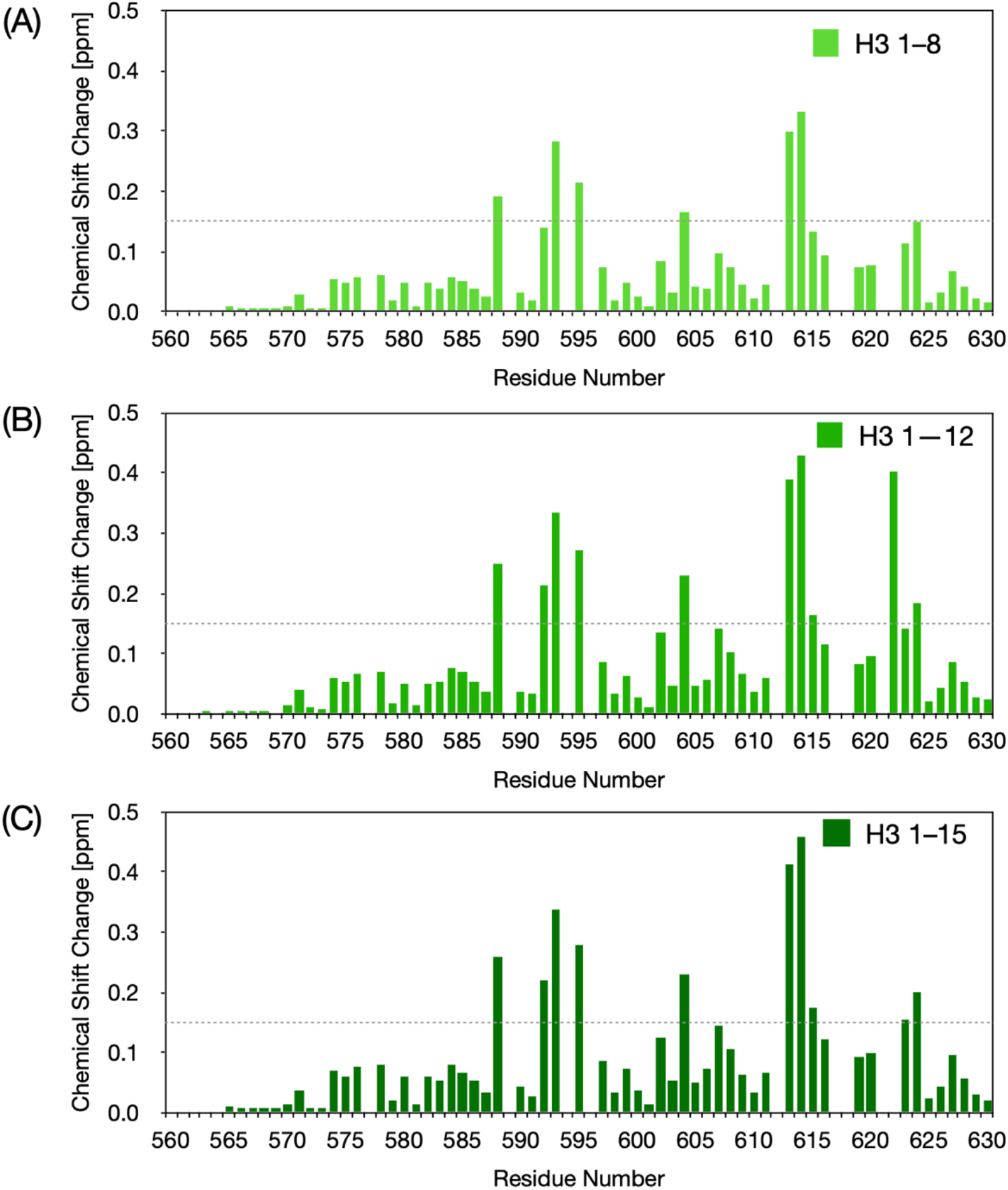
Magnitude of chemical shift changes observed upon addition of 250 µM histone H3 peptides of different lengths. Bar plots show the magnitude of chemical shift changes (ppm) as a function of KKT2 sequence for histone H3 peptides containing (A) 8, (B) 12, and (C) 15 residues. Note that Figure S6B is the same as Figure 7A, and is shown here for comparison. The dashed horizontal line at 0.15 ppm indicates the threshold used to identify significantly perturbed residues. For all plots, no chemical shift change is shown for I577, P589, P612, H617, K618 or Y621 due to the absence of peaks for these residues in the BEST-TROSY spectrum collected in the absence of peptide. In some plots, no chemical shift change is shown for G594, D596 and D622 because the peaks corresponding to these residues are still broadened beyond detection in the presence of 250 μM peptide indicating an interaction.

**Figure S7.**
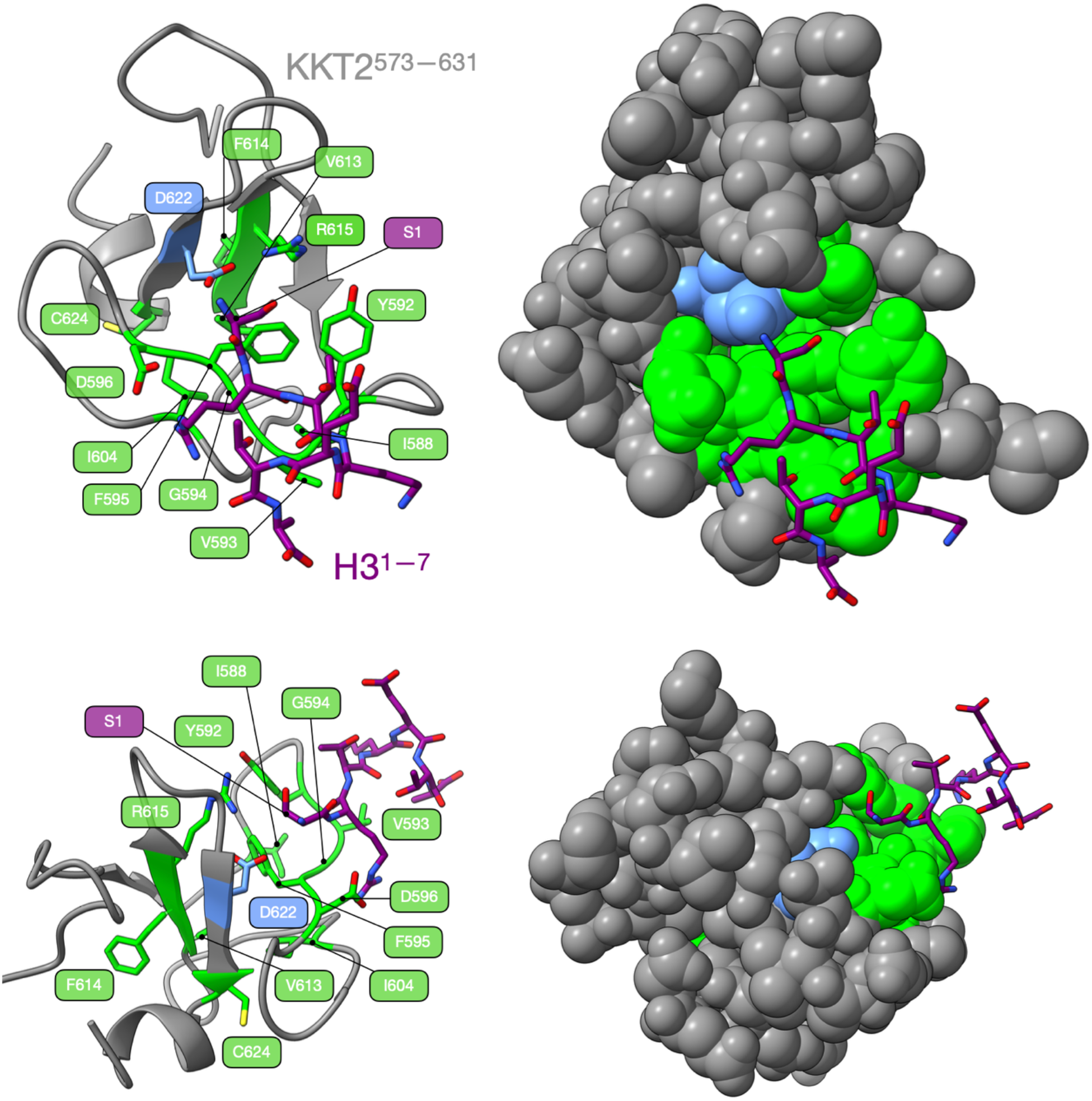
AF3 model of KKT2 CL domain with H3 peptide and residues that showed chemical shift changes upon H3 binding. Cartoon (left) and surface (right) representations are shown in two orientations. KKT2 is depicted in grey, and the H3 peptide is shown in purple sticks. Side chains of KKT2 residues exhibiting the most significant chemical shift perturbations (>0.15 ppm) are highlighted in lime green and displayed as sticks (left panel) and spheres (right panel). Residue D622 is highlighted in blue. Nitrogen, sulphur and oxygen atoms are labelled as red, yellow and blue, respectively.

**Figure S8.**
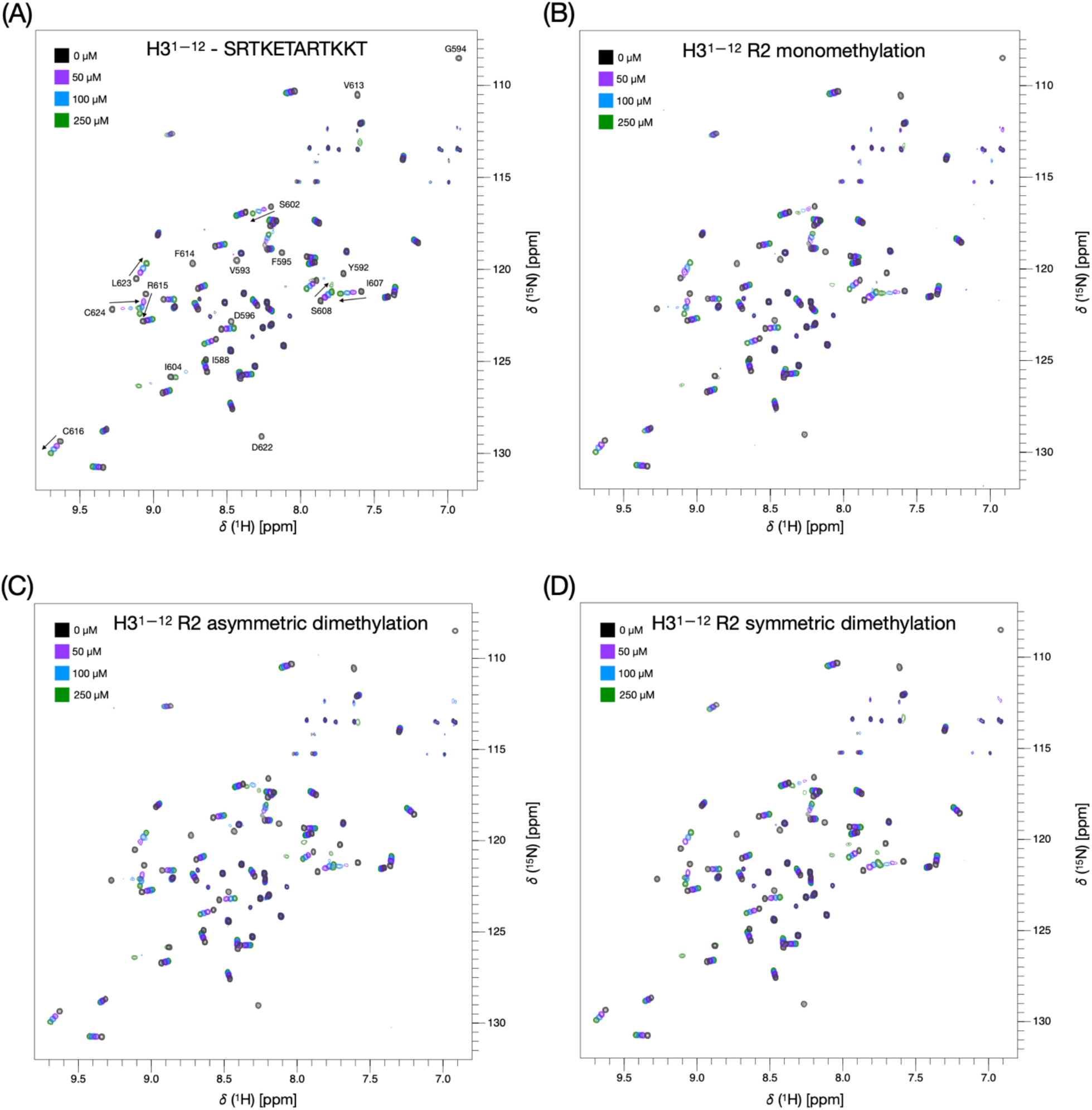
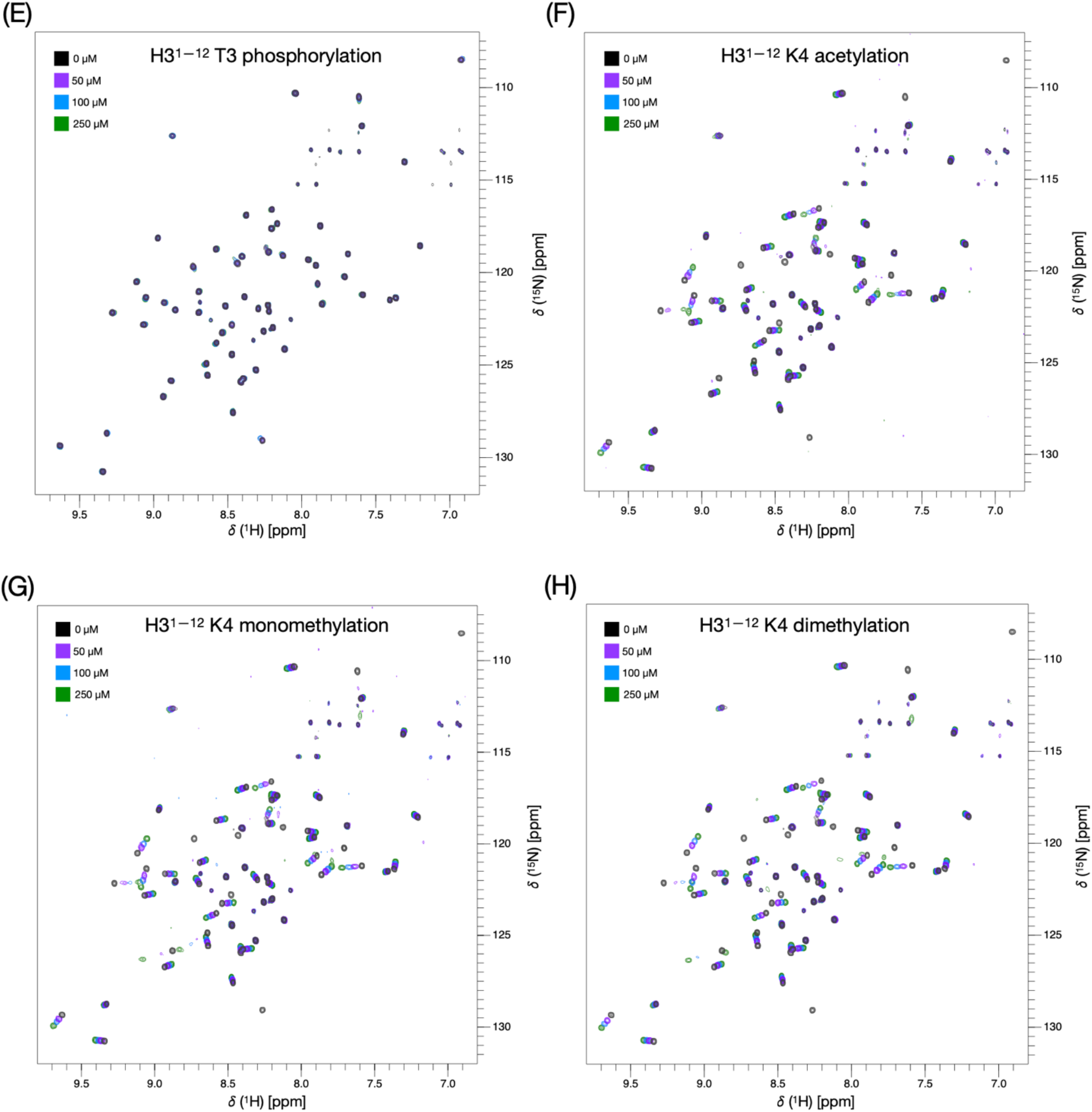
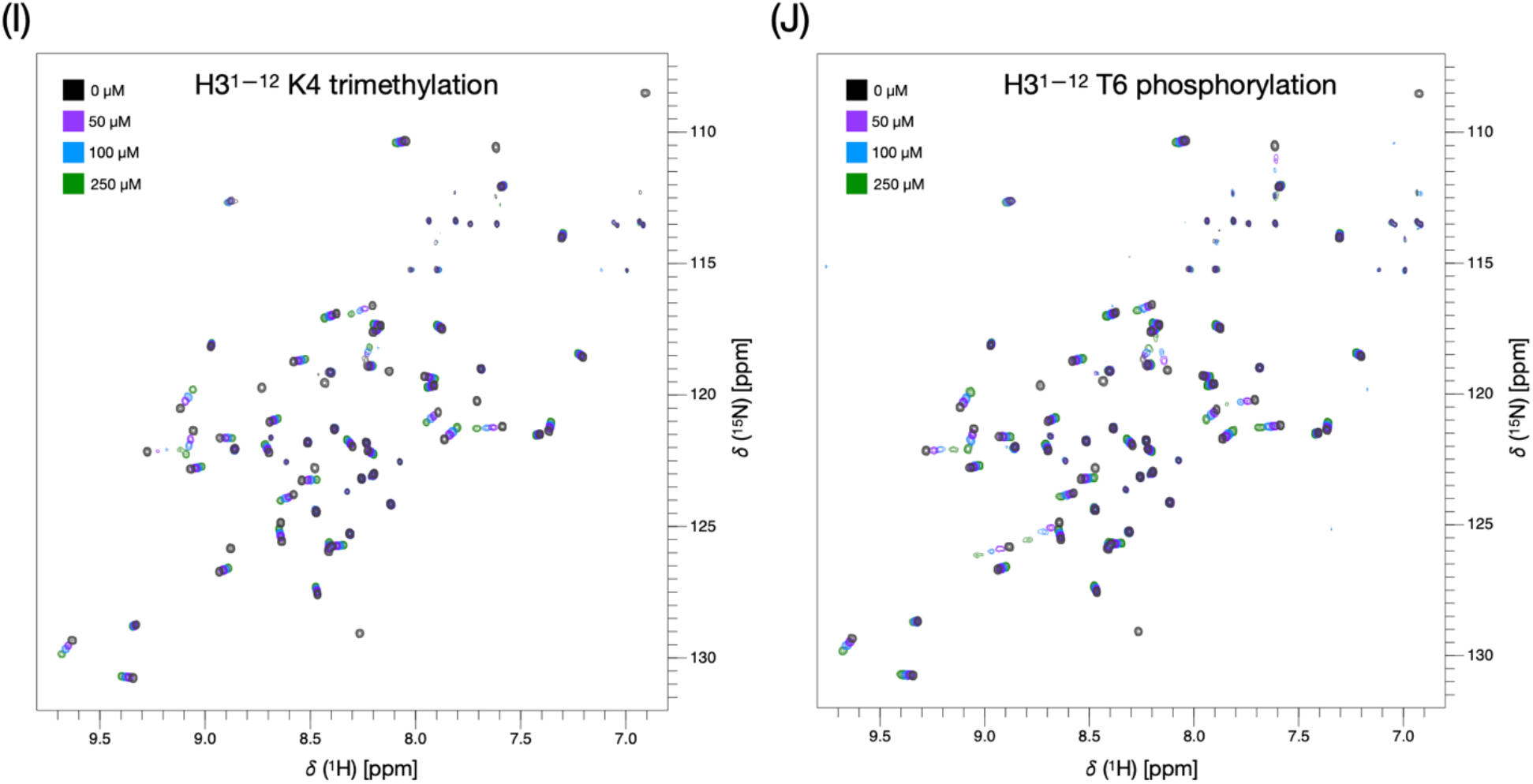
Effects of PTMs on H3 binding. Overlay of 750 MHz ^1^H-^15^N BEST-TROSY spectra of 50 μM KKT2 CL (KKT2^562–630^ in 25 mM HEPES, 100 mM NaCl and 0.5 mM TCEP at pH 7.2) in the absence (black) and with 50 (purple), 100 (blue) and 250 (green) μM of H3^1-12^ N-terminal peptides modified by (A) unmodified H3, (B) R2 monomethylation, (C) R2 asymmetric dimethylation, (D) R2 symmetric dimethylation, (E) T3 phosphorylation, (F) K4 acetylation, (G) K4 monomethylation, (H) K4 dimethylation, (I) K4 trimethylation, (J) T6 phosphorylation. Note that Figure S8A is the same as Figure 5A, and is shown here for comparison.

